# Role of human induced pluripotent stem cell-derived spinal cord astrocytes in the functional maturation of motor neurons in a multielectrode array system

**DOI:** 10.1101/614297

**Authors:** Arens Taga, Raha Dastgheyb, Christa Habela, Jessica Joseph, Jean-Philippe Richard, Sarah K. Gross, Giuseppe Lauria, Gabsang Lee, Norman Haughey, Nicholas J. Maragakis

**Affiliations:** Johns Hopkins University, Department of Neurology, Baltimore, MD, USA; Fondazione I.R.C.C.S. Istituto Neurologico Carlo Besta, Milan, Italy; Johns Hopkins University, Department of Neuroscience, Baltimore, MD, USA

**Keywords:** electrophysiology, spinal cord, glutamate receptor, gap junction, glia

## Abstract

The ability to generate human induced pluripotent stem cell (hiPSC)-derived neural cells displaying region-specific phenotypes is of particular interest for modeling central nervous system (CNS) biology *in vitro*. We describe a unique method by which spinal cord hiPSC-derived astrocytes (hiPSC-A) are cultured with spinal cord hiPSC-derived motor neurons (hiPSC-MN) in a multielectrode array (MEA) system to record electrophysiological activity over time. We show that hiPSC-A enhance hiPSC-MN electrophysiological maturation in a time-dependent fashion. The sequence of plating, density, and age in which hiPSC-As are co-cultured with MN, but not their respective hiPSC line origin, are factors that influence neuronal electrophysiology. When compared to co-culture with mouse primary spinal cord astrocytes, we observe an earlier and more robust electrophysiological maturation in the fully human cultures, suggesting that the human origin is relevant to the recapitulation of astrocyte/motor neuron cross-talk. Finally, we test pharmacological compounds on our MEA platform and observe changes in electrophysiological activity which confirm hiPSC-MN maturation. These findings are supported by immunocytochemistry and real time PCR studies in parallel cultures demonstrating human astrocyte mediated changes in the structural maturation and protein expression profiles of the neurons. Interestingly, this relationship is reciprocal and co-culture with neurons influences astrocyte maturation as well. Taken together these data indicate that in a human *in vitro* spinal cord culture system, astrocytes alter hiPSC-MN maturation in a time-dependent and species specific manner and suggest a closer approximation of *in vivo* conditions.

**Main Points:**

1. We developed a method for the co-culture of human iPSC-A/MN for multielectrode array recordings.
2. The morphological, molecular, pharmacological, and electrophysiological characterization of the co-cultures suggests bidirectional maturation.

## Introduction

Since the initial descriptions of techniques to generate human induced pluripotent stem cell-derived neurons (hiPSC-N) and astrocytes (hiPSC-A)^1^, there has been increasing interest in using these cells to recapitulate central nervous system (CNS) biology *in vitro* in both normal and disease states and to explore therapeutic strategies for neurological disorders^2^. With the increasing number of differentiation techniques, the electrophysiological characterization of hiPSC-N has become crucial to provide accurate measures of their function, beyond morphological studies, and, ideally, within an environment that resembles their *in vivo* counterparts.

Multi-electrode arrays (MEA) are particularly suited for these purposes^3^ as they enable the recording of large populations of neurons and their network activity, and have the potential to inform about neuron-glia interactions. This is achieved through the detection of extracellular voltages, which reflect the spike activity of local neuronal populations. The extracellular nature of the recordings allows for extended recordings that are particularly suitable for drug testing.

Previous MEA studies have focused on hiPSC-derived cortical neurons, either alone^4–6^ or in co-culture with primary rodent cortical astrocytes^7^ and more recently with hiPSC-derived astrocytes^5,8,9,10^. Despite an established body of evidence suggesting that astrocytes contribute to neuronal electrophysiological maturation by promoting synaptogenesis^11^, MEA- and human iPSC-based paradigms, still influenced by a traditionally neuron-centered perspective, have not yet been utilized to model this unique glial contribution. Recent literature^12^ has suggested that astrocytes are regionally heterogeneous and that astrocyte-neuron interactions may be influenced by their respective positional identities. However, hiPSC-A have been largely used as a homogenous and interchangeable cell type for MEA- and iPSC-based studies.

Here, for the first time, we describe a method in which the co-culture of spinal cord hiPSC-A and hiPSC-derived spinal cord motor neurons (hiPSC-MN) is optimized for obtaining MEA electrophysiological recordings. We examine the influences of hiPSC-A on the differentiation and electrophysiological maturation of hiPSC-MN, and demonstrate that hiPSC-A influence the capacity of hiPSC-MN to respond to neurotransmitters and their agonists/antagonists. Similarly, we show that hiPSC-A maturation is enhanced by the presence of hiPSC-MN in co-culture, thus allowing for a cross – talk that can be investigated with MEA recordings. This fully human *in vitro* platform for the electrophysiological evaluation of spinal cord astrocyte/MN interactions has the potential to more accurately model human diseases with spinal cord pathology, including spinal muscular atrophy (SMA) and amyotrophic lateral sclerosis (ALS).

## Methods

### Fibroblast collection and reprogramming

Induced pluripotent stem cells were reprogrammed from fibroblasts derived from skin biopsies under Johns Hopkins IRB protocols NA_00021979 and NA_00033726. Consents included the authorization to use DNA, fibroblasts, or cell lines derived from fibroblasts for research purposes only. Fibroblasts were cultured and induced pluripotent stem cell lines were created and initially characterized by one of the authors (G.Lee) and with an NIH--‐sponsored commercial agreement with iPierian (USA), who used a 4-vector method as we have previously described^13^. For this study, we utilized two different cell lines from patients with no known disease diagnosed at time of fibroblast collection: CIPS and GM01582.

### iPSC culture, hiPSC-MN and hiPSC-A differentiation

Induced pluripotent stem cells were maintained in E8™ medium with Essential 8™ supplement (Gibco-Thermo Fisher Scientific) and passaged once every 5-6 days using Dispase (StemCell Technology). To maintain pluripotency and limit spontaneous differentiation, the stem cell colonies were manually cleaned before passaging.

Induced pluripotent stem cells were differentiated into spinal cord neural progenitor cells (NPCs) following modifications of a 25-30 day protocol described previously^14^ that relies on extrinsic morphogenetic instructions to pattern NPCs along the rostro-caudal and dorso-ventral axis, and includes: neuralization, through dual SMAD signaling inhibition, using LDN193189 (Stemgent) and SB431542 (Millipore Sigma), followed by caudalization and ventralization, through retinoic acid (RA) and purmorphamine (PMN) (Millipore Sigma) (Figure 1A).

**Figure 1.**
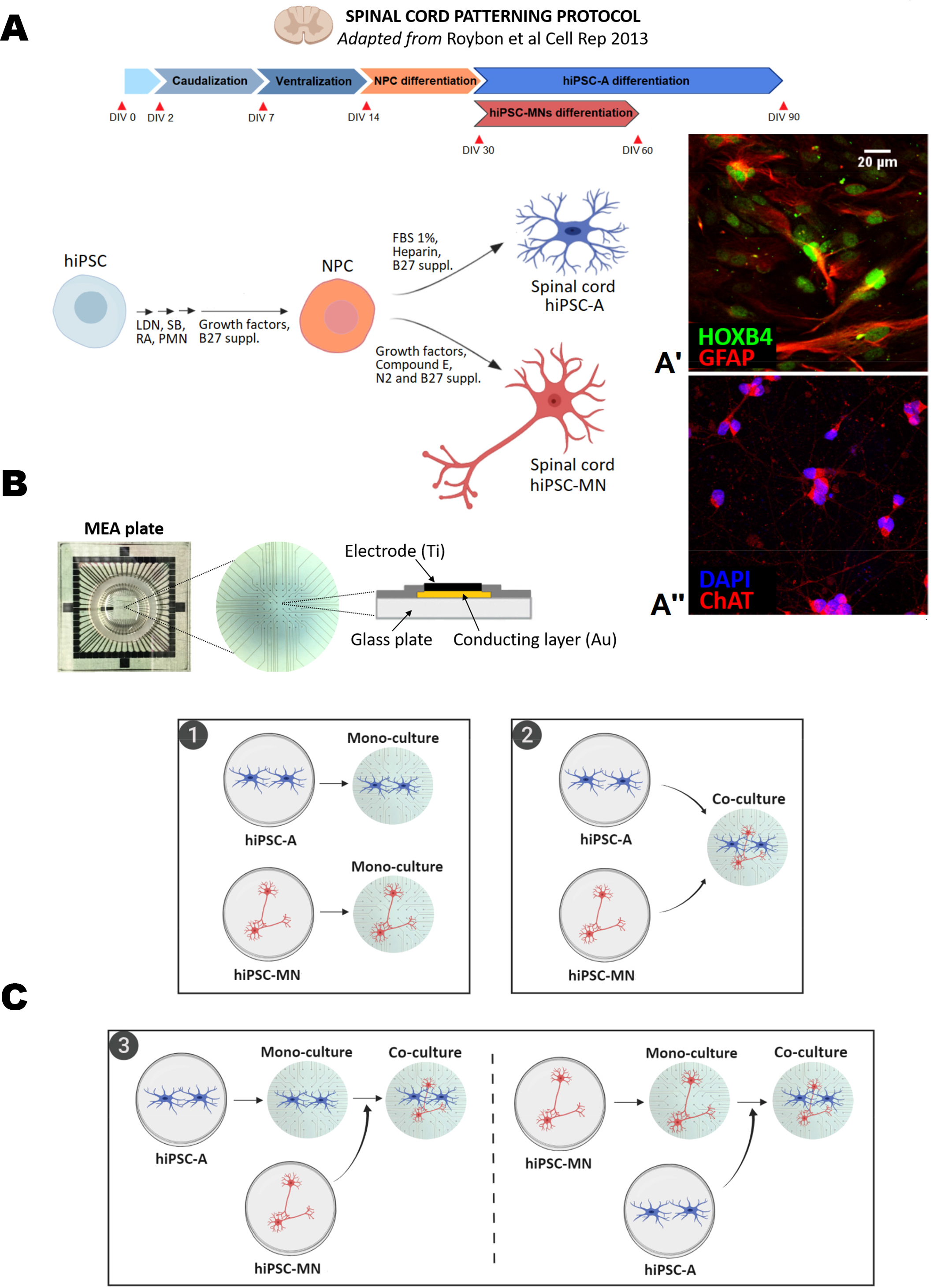
**A: Schematic illustration of the differentiation protocol to generate spinal cord hiPSC-A and hiPSC-MN.** Two representative immunofluorescence images of positional identity markers HOXB4 (***A’***), a transcription factor involved in spinal cord development, and ChAT (***A”***), an enzyme responsible for the biosynthesis of acetylcholine in spinal cord motor neurons. **B**: Photograph of MEA plate, with detail on its set of 60 microelectrodes and schematic illustration of an individual electrode. **C**: Diagram of human hiPSC-A/-MN cultures on MEA plate: *(**1**)* hiPSC-A or –MN mono-cultures, where astrocytes or neurons were cultured alone, *(**2**)* simultaneous co-culture, where both cell types were mixed and plated simultaneously, and *(**3**)* serial co-cultures, where hiPSC-A (*left*) or hiPSC-MN (*right*) where plated alone prior to the addition of neurons and astrocyte, respectively. Abbreviations: LDN, LDN193189; SB, SB431542; RA, retinoic acid; PMN, purmorphamine; FBS, fetal bovine serum.

For motor neuron differentiation, NPCs were cultured with “neuronal differentiating medium”, comprised of Neurobasal (Gibco-Thermo Fisher Scientific), enriched with N2 and B27 supplement (Gibco-Thermo Fisher Scientific), L-glutamine, non-essential amino acids (Gibco-Thermo Fisher Scientific), penicillin/streptomycin and supplemented with growth factors (RA, PMN, ascorbic acid, recombinant human-brain-derived neurotrophic factor, glial cell line-derived neurotrophic, insulin-like growth factor 1, ciliary neurotrophic factor). The addition of Compound E (Santa Cruz Biotechnology), a γ-secretase inhibitor, enhanced neuronal differentiation into motor neurons. To prevent astrocyte over-proliferation, neuronal cultures were treated once with 0.02μM cytosine arabinoside (ara-C) (Millipore Sigma) for 48 h. The medium was then changed every other day. At 60 days in vitro (DIV), this protocol has been shown to generate a population of spinal cord neurons, with a majority of neurons expressing MN markers, including choline acetyltransferase (ChAT)^14^ (Figure 1A”) and ISL1/2.

For the generation of spinal cord hiPSC-A (Figure 1A), NPCs were cultured with “astrocyte differentiating medium”, containing DMEM:F12 (Gibco-Thermo Fisher Scientific), enriched with B27 supplement (Gibco-Thermo Fisher Scientific), L-glutamine, non-essential amino acids, penicillin/streptomycin, heparin (Millipore sigma), and supplemented with 1% fetal bovine serum (FBS) (Gibco-Thermo Fisher Scientific). The medium was changed every other day, and cultures were passaged when confluent. After 90 DIV, this protocol generates a population of mature astrocytes^14,15^ which express HOXB4, a marker of spinal cord regional identity^14^ (Figure 1A’).

### Mouse primary spinal cord astrocyte isolation and culture

The isolation and culture of spinal cord astrocytes from postnatal mouse spinal cord was performed as previously published^16^. Briefly, the spinal cord was dissected from post-natal day 2 (P2) mouse pups, and the meninges removed to avoid the subsequent contamination of the astrocyte culture with fibroblasts. The tissue was then enzymatically dissociated with papaine (Worthington Biochemical) and deoxyribonuclease I (Sigma), and mechanically triturated to generate a single cell suspension. Cells were plated on poly-L-lysine-coated T25 flasks, in a medium containing DMEM with 10% FBS and 1% penicillin–streptomycin. Cells were allowed to recover and proliferate for 2 weeks prior to plating.

### Multi-electrode array culture

Multi-electrode array plates (60MEA200/30iR-Ti-gr, MultiChannel Systems; MCS) were used for the recording of hiPSC-MN cultures (Figure 1B). Prior to plating, the glass surface of the MEA plates was first treated with 1% Terg-a-zyme (Sigma Aldrich) overnight at room temperature, and then with O_2_ plasma (Harrick Plasma) for 1 minute. The plates were then coated with poly-ornithine (Millipore Sigma; concentration: 100μg/mL) and laminin (Thermo Fisher Scientific; concentration: 5μg/ml). Recordings were conducted using an MEA2100 System (MCS) on a stage heated to 37 °C.

For MEA recording, hiPSC-MN, derived from the GM01582 cell line, and hiPSC-A, derived from the CIPS cell line, were cultured as follows (Figure 1C). For hiPSC-MN mono-cultures, neurons were plated after 60DIV at a density of 5×10^5^ cells/plate; this neuron density was maintained as a constant for all culture conditions. For hiPSC-A mono-cultures, astrocytes were plated after 90DIV at a density of 1×10^5^ cells/plate. For the astrocyte first serial co-culture, we first plated hiPSC-A cultured for 86DIV at a density of 1×10^5^ cells/plate; four days later, we added hiPSC-MN cultured for 60DIV. For the neuron first serial co-culture, we first plated hiPSC-MN cultured for 56DIV; four days later, we added hiPSC-A cultured for 90DIV and then plated at a density of 1×10^5^ cells/plate. For the simultaneous co-culture of hiPSC-A/hiPSC-MN, both cell types were mixed and plated simultaneously on MEA plates at the densities noted above, after 90DIV and 60DIV, respectively.

As variations of the above-mentioned simultaneous co-culture, we also used the following: hiPSC-A which had been cultured for 60DIV (“immature hiPSC-A”) instead of 90DIV, a lower density of hiPSC-A (i.e. 0.5×10^5^ cells/plate instead of 1×10^5^ /plate), and hiPSC-A differentiated from same iPSC line (“isoclonal”) as hiPSC-MN (i.e. both lines were GM01582), instead of using different control iPSC lines for astrocytes (CIPS) and neurons (GM01582) (i.e. “heteroclonal”). Finally, we used primary mouse spinal cord astrocyte (“mouse A”), instead of human iPSC-A, simultaneously co-cultured with hiPSC-MN for some studies.

For all MEA culture conditions, the culture medium was changed at day 1 of MEA plating to “neuronal” medium enriched with 5% FBS, 0.5 μg/ml laminin and 2.0 μg/ml Amphotericin B (Gibco). Cells were fed with a half medium exchange every 3 days.

### Multielectrode array recordings

The MEA plates used for this study have 60 electrodes, including 59 active and 1 inactive that represents the reference for unipolar acquisition. Voltage measurements were made at a sampling rate of 25 kHz/channel using MC_RACK software (MCS) and filtered using a second order butterworth filter with a 200Hz cutoff frequency. Spikes were identified as instantaneous time points of voltages that exceed a threshold of at least five standard deviations below baseline. Bursts were defined as activity with >4 spikes/0.1 sec. Hypersynchronous (or “network”) bursts were defined when over 60% of active electrodes fired within one 20-ms bin, as previously described ^17^. Spike trains were exported into an HDF5 format and further analyzed using MEAnalyzer^18^. The following electrophysiological parameters were analyzed: spike and burst rate, percentage of spiking and bursting electrodes (on the overall 59 active electrodes). Functional connectivity graphs based on spike rate were generated using MEAnalyzer as recently described^19^. The recording of the baseline activity of MEA plates was performed weekly over one a minute period, for 4 weeks after plating (i.e. 4 time points, week 1, 2, 3, and 4).

For pharmacological testing, we recorded: baseline MEA activity for 1 minute, activity 1 minute after 100 μL “fresh” medium exchange with the drug vehicle, and 1 to 10 minutes after the exchange of 100 μL of medium containing the compound of interest and the vehicle. We tested the following compounds targeting ion channels and/or neurotransmitter receptors, based on published literature^7^: 100 mM potassium chloride (KCl); 5μM of the α-amino-3-hydroxy-5-methyl-4-isoxazolepropionic acid (AMPA)/kainate receptor agonist, kainic acid (KA) (Abcam); 50μM of the AMPA/kainite receptor antagonist, cyanquixaline (CNQX) (Sigma Aldrich); 10 μM of the gamma amino butyric acid (GABA) receptor antagonist, bicuculline (Sigma Aldrich). We also tested compounds targeting glia, and verified their effect on neuronal firing; these included: GAP19 (Tocris), a connexin 43 hemichannel blocker^20^, at concentrations of 34 μM and 340 μM and dihydrokainic acid (DHK) (Tocris), an excitatory amino acid 2 (EAAT2) selective blocker, at concentrations^21,22^ of 50 μM and 300 μM. We analyzed the effects of the above-mentioned compounds on 3 MEA plates at two different time points and when at least one electrode showed spike rate > 0: at week 1 or 2 of culture, and at week 3 or 4.

### Immunocytochemistry

In parallel experiments with MEA and using the same cell densities, neurons and astrocytes were plated in 24 well plates on glass cover-slips for immunocytochemistry. Two time points were considered: 1 week and 4 weeks of mono- or simultaneous co-culture.

Cells were fixed with 4% paraformaldehyde for 10 minutes and then washed with phosphate-buffered saline (PBS) three times. The cells were then permeabilized with 0.1% Triton™ X-100 (Millipore Sigma) in PBS for 10 minutes and washed with PBS three times. A blocking solution with 3% bovine serum albumin (BSA) in PBS was then applied for 1 hour. Coverslips were stained with primary antibodies in a blocking solution containing 3% BSA in PBS and 3% species specific serum and incubated overnight at 4ºC. The next day, cells were washed with 3% BSA in PBS three times and incubated with appropriate secondary antibodies (Thermo Fisher Scientific, Alexa Fluor Dyes; concentration: 1:1000) and Hoechst® (Thermo Fisher Scientific; concentration: 2 μg/ml) in a blocking solution with 3% BSA in PBS and 3% species-specific serum, for 1 h at room temperature. Finally, the coverslips were washed with 3% BSA in PBS three times and mounted with Prolong gold with DAPI® (Thermo Fisher Scientific) and stored until ready to image. The primary antibodies used for this study are listed in Table 1.

**Table 1.**
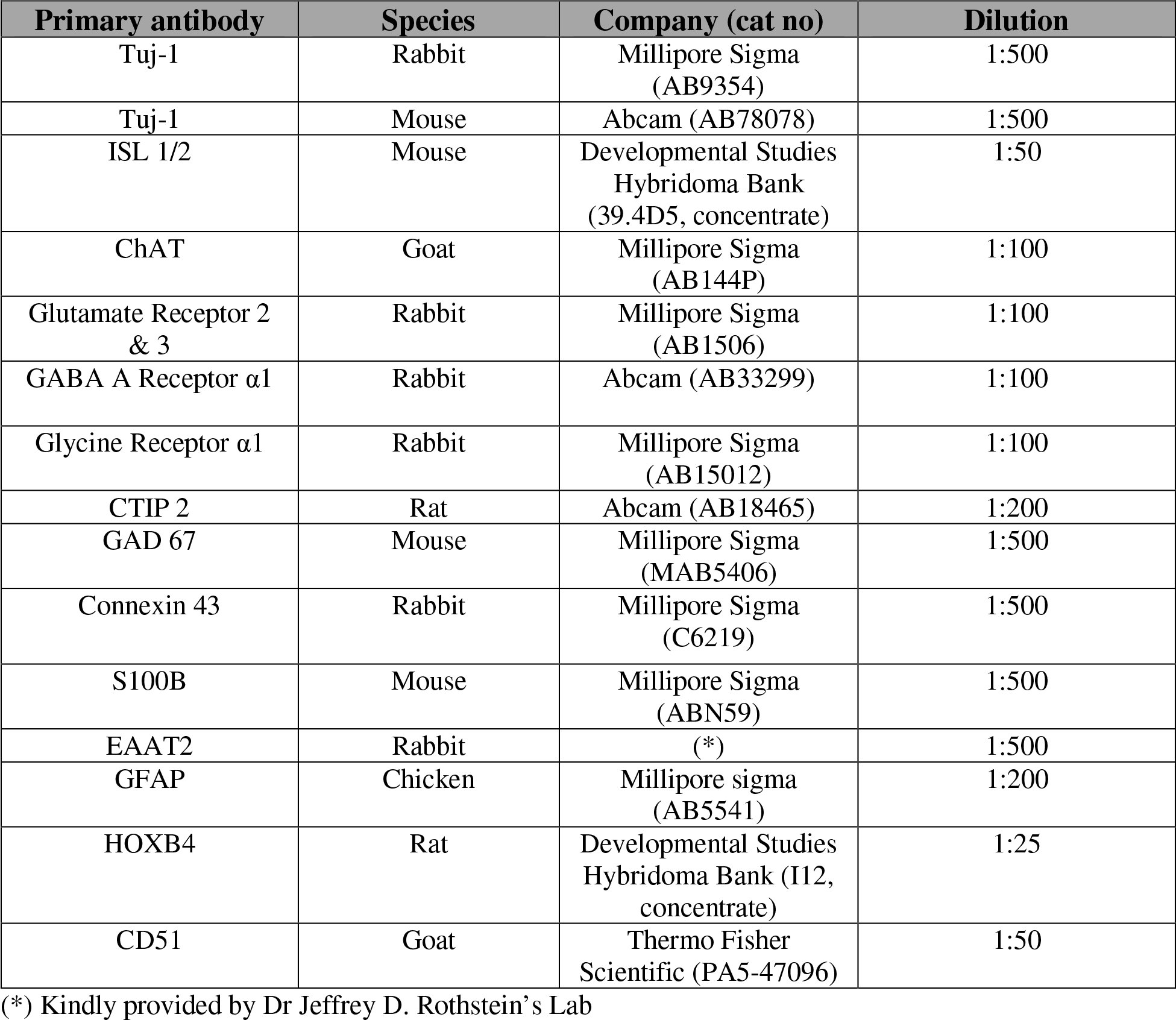
List of primary antibodies used in this study.

| **Primary antibody** | **Species** | **Company (cat no)** | **Dilution** |
| --- | --- | --- | --- |
| Tuj-1 | Rabbit | Millipore Sigma (AB9354) | 1:500 |
| Tuj-1 | Mouse | Abcam (AB78078) | 1:500 |
| ISL 1/2 | Mouse | Developmental Studies Hybridoma Bank (39.4D5, concentrate) | 1:50 |
| ChAT | Goat | Millipore Sigma (AB144P) | 1:100 |
| Glutamate Receptor 2 & 3 | Rabbit | Millipore Sigma (AB1506) | 1:100 |
| GABA A Receptor α1 | Rabbit | Abcam (AB33299) | 1:100 |
| Glycine Receptor α1 | Rabbit | Millipore Sigma (AB15012) | 1:100 |
| CTIP 2 | Rat | Abcam (AB18465) | 1:200 |
| GAD 67 | Mouse | Millipore Sigma (MAB5406) | 1:500 |
| Connexin 43 | Rabbit | Millipore Sigma (C6219) | 1:500 |
| S100B | Mouse | Millipore Sigma (ABN59) | 1:500 |
| EAAT2 | Rabbit | (*) | 1:500 |
| GFAP | Chicken | Millipore sigma (AB5541) | 1:200 |
| HOXB4 | Rat | Developmental Studies Hybridoma Bank (I12, concentrate) | 1:25 |
| CD51 | Goat | Thermo Fisher Scientific (PA5-47096) | 1:50 |

(\*) Kindly provided by Dr Jeffrey D. Rothstein's Lab

Images were acquired on a Zeiss fluorescence microscope (Zeiss ApoTome.2), using 20x and 63x oil magnification and analyzed using Image J software (NIH) and Fiji package of Image J. Five images were obtained for each coverslip, and 3 coverslips were utilized for each condition. Cell counts were performed by a person who was blinded to the experimental conditions.

### Quantitative analysis of neurites and synapses

For the morphological analysis of neurons, we used TUJ1 immunostaining and 63x oil magnification images. We manually traced the diameter of individual neurites, and for each neuron, we defined the primary neurite as the largest projection from the cell body. The area of each neuronal soma was traced manually. For these analyses, 50 randomly selected neurons were considered for each condition.

To define the complexity of neuronal connections, we first tracked individual neurites on 63x TUJ1 images using the simple neurite tracer software plugin on the Fiji package of image J^23^ and then performed a Sholl analysis^24^ on the neurite mask, with soma-centered concentric circles of increasing radius (10 μm increment); for analysis purposes, we considered the mean number of intersection for each individual neuron.

Finally, we quantified synapses as the number of co-localized punctae of synaptophysin and PSD-95 antibody staining on 63x oil images, as previously described^25^. This immunocytochemistry-based protocol determines the mean number of co-localized punctae within a defined region of interest (ROI) surrounding neuronal soma. We used circular regions, one-cell diameter radially around the soma of interest, which was identified with TUJ1 staining. For Sholl analysis and for the quantification of synapses, we considered a minimum of n=10 neurons/per coverslip, randomly selected, and 3 coverslips for each condition.

### qPCR of neuronal and astrocyte RNA transcripts

Human-specific primers for neuronal and glial target genes (Table 2) were designed, and the primer sequences were confirmed by BLAST analysis and tested for specificity. Universal eukaryotic 18S rRNA primers were used as endogenous control, allowing for comparisons between human-human and human-mouse co-cultures, as previously published ^26,27^.

**Table 2.**
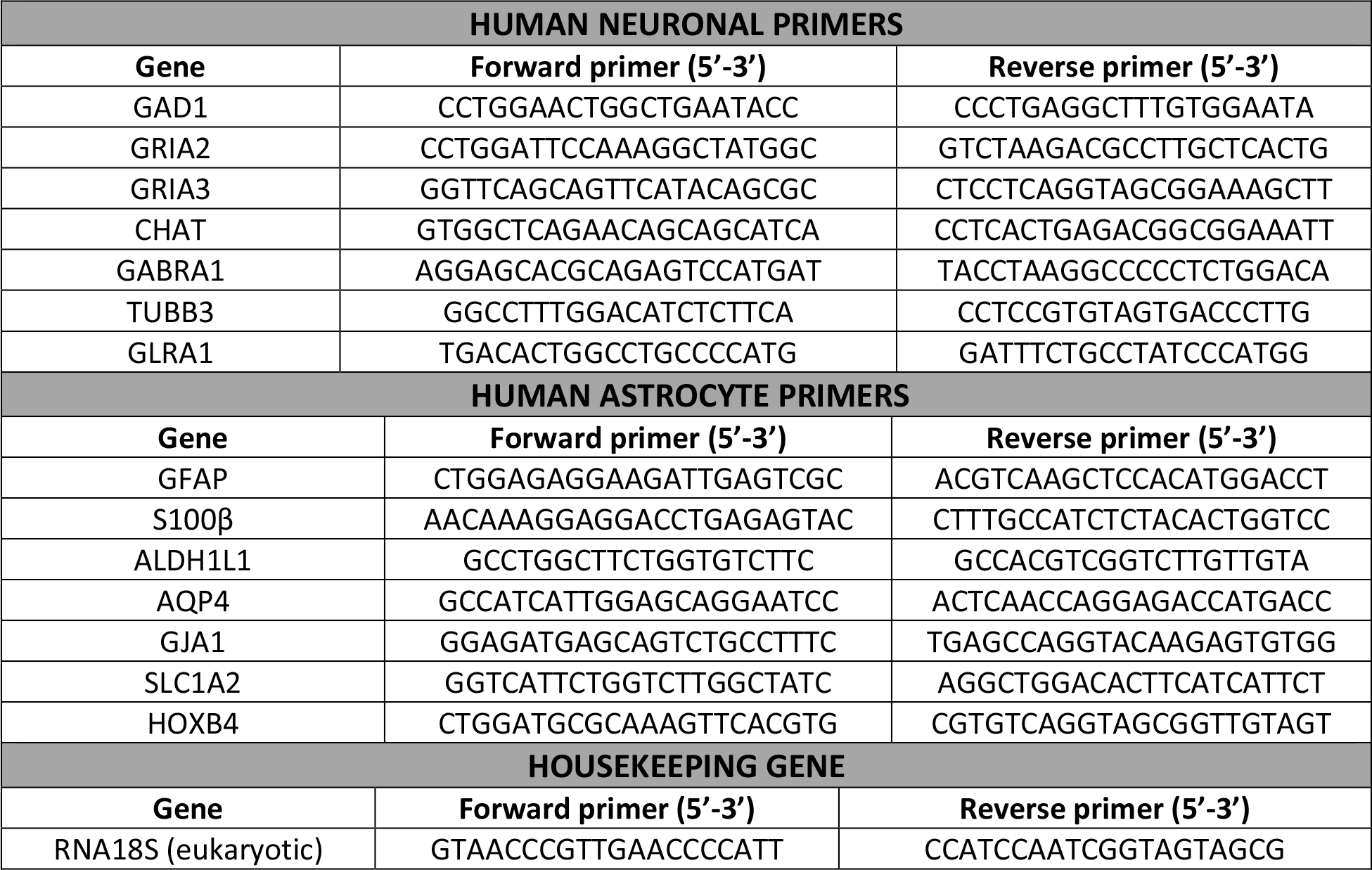
List of primers used for RT-PCR. Gene nomenclature follows the indications of the HUGO Gene Nomenclature Committee (https://www.genenames.org).

Human iPSC-A, immature hiPSC-A, mouse primary spinal cord astrocytes and hiPSC-MN were grown as monolayers in 6 well plates (3 wells per condition), in the same conditions, densities and time points used for immunostaining and in parallel experiments. After 1 and 4 weeks *in vitro*, cell cultures were lysed and homogenized using TRIzol^TM^ reagent (Invitrogen). Human iPSC-A and hiPSC-MN from mono-cultures were collected together and in the same volume of TRIzol (i.e. 0.3 ml/well) used for simultaneous co-cultures. With this strategy, and since the number of TUJ1^+^/DAPI^+^ and TUJ1^−^/DAPI^+^ cells was not significantly different between monocultures and co-cultures (as showed in the results section), we anticipated comparable loading controls from the initial samples.

Total RNA was then extracted using RNeasy Mini Kit (Qiagen, 74104). The quality and quantity of purified RNA was examined using a NanoDrop spectrophotometer (Thermo Fisher Scientific). The RNA was converted into cDNA using the iScriptTM cDNA synthesis kit (Bio-Rad) following manufacturer's instructions. The cDNA was then amplified using the Fast SYBR green PCR master mix (Applied Biosystems) in technical duplicates for each individual sample. The expression of target genes was normalized to 18s levels using the comparative CT method ^28^.

### Data presentation and statistical Analysis

All data were analyzed using Graph Pad Prism software (La Jolla, CA). Data are presented as bar graphs for mean ± SEM, with individual observations visualized as scatter plot, or as box (interquartile range and median) and whiskers (minimum and maximum). Data were analyzed using one-way ANOVA, followed by Tukey’s test for multiple comparisons and a two-tailed unpaired t-test, as appropriate. Data distribution was assumed to be normal but this was not tested. The statistical significance was set at *p*<0.05. Unless stated otherwise, all experiments were performed in technical triplicates.

For qPCR analyses, the ΔΔCT values were normalized to mono-cultures (i.e. “hiPSC-A” or “hiPSC-MN), to account for the effect of co-cultures.

For the analysis of MEA baseline activity, we considered the active electrodes only (i.e. electrodes with spike rate > 0 using MEAnalyzer^18^), and their mean activity as a measure of the overall electrophysiological activity of the cell culture. To account for variability in the culturing process, analyses were performed only on parallel plating.

In order to define the effect of KCl and neurotransmitter modulators on MEA activity, we recorded for 3 minutes, and compared the mean activity at baseline (1 minute), after medium with vehicle exchange (1 minute) and after drug addition (in the same volume of medium and with the same concentration of vehicle) (1 minute). For GAP19 we recorded for 3 minutes after drug addition, since changes in MEA activity were slightly slower. For DHK we recorded for 10 minutes after drug addition, to detect even slower effects; since changes in the pH in the medium could occur at atmospheric conditions for longer recordings, we compared the activity after DHK addition to parallel plates where we evaluated changes up to 10 minutes after the addition of the vehicle. All experiments were performed in experimental triplicates.

## Results

### hiPSC-A influence hiPSC-MN maturation

In order to develop a reliable and reproducible platform to study the influences of hiPSC-A on hiPSC-MN electrophysiology, we first sought to optimize the cell culturing protocol on MEA plates, and to determine the optimal sequence of plating hiPSC-A and hiPSC-MN (Figure 2). Human iPSC-A, after proper treatment of the plate surface and at the densities noted above, generated a confluent monolayer (Figure 2A, **box 1).** The culture of hiPSC-MN alone resulted in large aggregates of neurons, which were arranged over a few electrodes (Figure 2A**, box 2 and** Figure 2B). The culture of hiPSC-A into a monolayer followed by plating hiPSC-MN resulted in fewer large neuronal aggregates and a more even distribution of cells amongst the electrodes (Figure 2A**, box 3**). Plating hiPSC-MN followed by the addition of hiPSC-A had similar results (not shown). Finally, we found that the simultaneous culture of hiPSC-A and hiPSC-MN resulted in a more evenly distributed populations of MN with fewer aggregates of MN and fewer numbers of cells in each aggregate (Figure 2A, **box 4 and** Figure 2B). These observations were consistent among 15 different MEA cultures.

**Figure 2.**
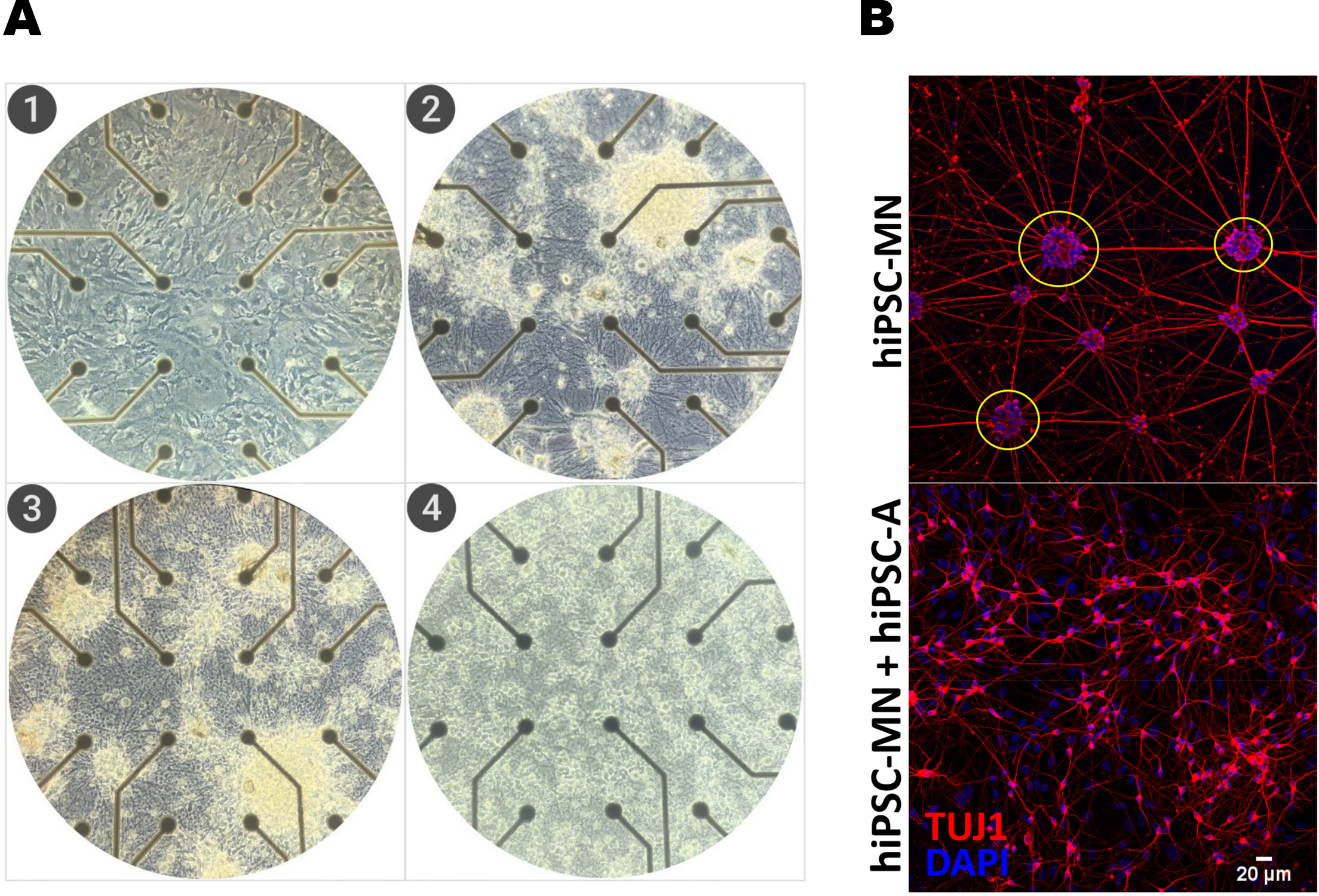
**A**: **Phase contrast images of hiPSC-A/-MN cultures on MEA plates (20x magnification).** The following culture conditions were considered: hiPSC-A alone *(**1**)*, hiPSC-MN alone *(**2**)*, serial (astrocytes first serial co-culture is shown) *(**3**)* and simultaneous *(**4**)* co-culture. **B**: **Immunofluorescent images (20x magnification).** TUJ1 immunostaining defines neuronal populations. Neuronal clusters (yellow circle) are more evident in hiPSC-MN monocultures than in hiPSC-MN and hiPSC-A simultaneous co-culture.

We then asked whether the presence of astrocytes and the time in culture would result in a change in neuronal subtype composition and maturation. When hiPSC-MN were cultured in mono-cultures for 1 week, we found that >90% of TUJ1^+^ neurons had a motor neuron identity, as demonstrated by ChAT immunoreactivity (Supplementary Figure 1A). This percentage was not significantly different when compared to astrocyte-neuron co-cultures (93.1 ± 3.0 % vs 92.1 ± 4.2 %, respectively) suggesting that motor neuron identity is defined early during iPSC differentiation in spinal cord neural progenitors. This result was confirmed by ISL-1/2 staining of a subset of MN^29^, which was for both conditions between 25-30% of TUJ1^+^ cells (28.4±0.8% vs 27.4±1.5%, respectively). Consistent with the spinal cord identity of our neuronal culture conditions, less than 2% of our neuronal population was positive for CTIP2 (a marker of corticospinal motor neuron identity)^30^. Beside ChAT^+^ motor neurons, neuronal cultures showed a small proportion (<5%) of GABAergic interneurons, as suggested^31^ by the immunoreactivity to GAD67. These percentages were not influenced by the time in culture, as they were not significantly different after 4 weeks of mono- or co-culture, thus reinforcing previous observations^32^ that spinal cord motor neuron fate is determined early during hiPSC differentiation (Supplementary Figure 1A).

The immunoreactivity for neuronal neurotransmitter receptors (Figure 3), including AMPA, GABA, and glycine were all significantly increased by the presence of hiPSC-A, both after 1 week (glutamate 2/3 receptor: 7.1±0.6% vs 28.9±0.9%, *p*<0.001; GABA receptor: 6.6±1.1% vs 16.4±0.9%, *p*<0.001; glycine receptor: 6.2±1.3% vs 10.9±2.0, p<0.05) and 4 weeks (glutamate 2/3 receptor: 7.3±0.4% vs 28.3±2.5%, p<0.001; GABA receptor: 4.8±1.4% vs 17.4±1.9%, p<0.001; glycine receptor: 5.1±0.4% vs 10.2±1.6, p<0.05) of co-culture. In parallel to neurotransmitter immunoreactivity, we found that the number of synapses, as demonstrated by PSD95 / synaptophysin staining was significantly increased when hiPSC-MN were co-cultured with hiPSC-A (Figure 3). The number of synapses increased over time with the culture of hiPSC-MN alone (6.9±0.8 vs 54.2±8.5, *p*<0.001), but to a greater degree (25.5±2.3 vs 85.1±5.1, *p*<0.01) in the presence of astrocytes (week 1 mono-vs co-culture, *p* < 0.01; week 4 mono-vs co-culture, *p* <0.001) (Figure 3).

**Figure 3.**
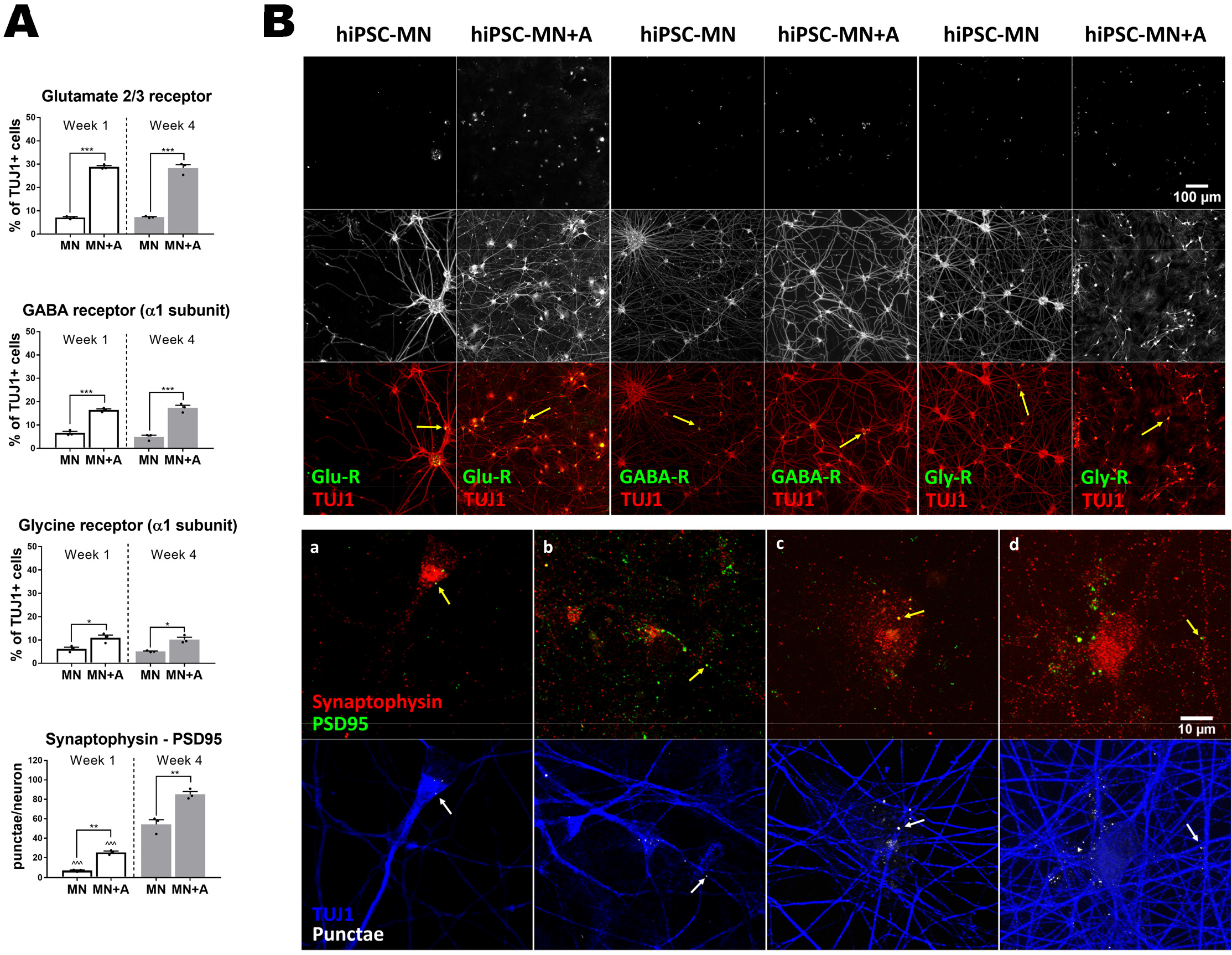
**A**: **Immunofluorescent quantification of neurotransmitter receptors and synapses.** Neurotransmitter receptors (glutamate, GABA and glycine) were quantified as percentage of TUJ1 immunopositive cells (mean of *n*=3 coverslips per culture condition and time-point). Synapses were counted as co-localized punctae of synaptophysin, a pre-synaptic marker, and PSD95, a post-synaptic marker, per individual neuron (mean of *n*=3 coverslips per condition, with a minimum of 10 neurons per coverslip). hiPSC-MN mono-cultures (MN) *vs* hiPSC-MN + hiPSC-A co-cultures (MN+A) were compared within each time-point after plating (*), i.e. week 1 (white bar) and week 4 (grey bar). Similarly, time-point observations (week 1 *vs* week 4) were compared within each condition (^). * *or* ^ *p*<*0.05;* ** *or* ^^ *p*<*0.01;* *** *or* ^^^ *p*<*0.001*. **B**: **Representative immunohistochemical images for neurotransmitter receptors and synapses. *Top***: Immunoreactivity for receptors, i.e. glutamate 2/3 receptor, GABA A receptor and glycine receptor in hiPSC-MN+A co-cultures compared to hiPSC-MN monocultures. Representative images were taken 1 week after plating. ***Bottom***: Co-localized synapthophysin/PSD95 punctae per neuron in hiPSC-MN+A co-cultures (***b,d***) compared to mono-cultures ***(a,c)***, from week 1 ***(a,b)*** to week 4 ***(c,d)***. Abbreviations: Glu-R, glutamate 2/3 receptor; GABA-R, GABA A receptor; Gly-R, glycine receptor.

We also investigated motor neuron maturation as defined by morphological parameters outlined in Figure 4. First, we considered *n*=100 randomly selected hiPSC-MN neurites in mono- or co-cultures, and found that the mean value of the diameter distribution was significantly different among conditions: 1.5±0.6μm vs 2.1±0.7μm for week 1 motor neurons in mono-vs co-culture, respectively (*p*<0.001), and 2.2±0.6μm vs 3.0±1.7μm for week 4 mono-vs co-cultures, respectively (*p*<0.001). Notably, after 4 weeks of co-culture, a subgroup of larger diameter neurites emerged, as seen in the histogram in Figure 4A. We then considered the largest neuronal process of *n*=50 randomly selected neurons, and found that increased overtime, with slightly higher values in the presence of hiPSC-A though not statistically significant: 5.4±1.8μm vs 9.1±3.1μm, *p* <0.001 for monocultures, and 5.9μm ±2.7μm vs 9.7μm ±3.9μm, p<0.001 for co-cultures. The area of the neuronal cell body (*n*=50 neurons) significantly increased over time in hiPSC-MN mono-cultures (125.9±28.9μm^2^ vs 203.9±32.6μm^2^, *p*<0.001), and more robustly (*p*<0.001) in the presence of hiPSC-A (131.1±26.1μm^2^ vs 230.9±26.3μm^2^, *p*<0.001). Finally, to quantify the complexity of connections between neurons we performed a Sholl analysis and found that the mean number of intersections per neuron increased significantly overtime (2.2±0.3 vs 14.5±0.9, *p*<0.05), particularly (*p*<0.001) when hiPSC-MN where in co-culture with hiPSC-A (1.9±0.2 vs 38.6±8.3, *p*<0.001).

**Figure 4.**
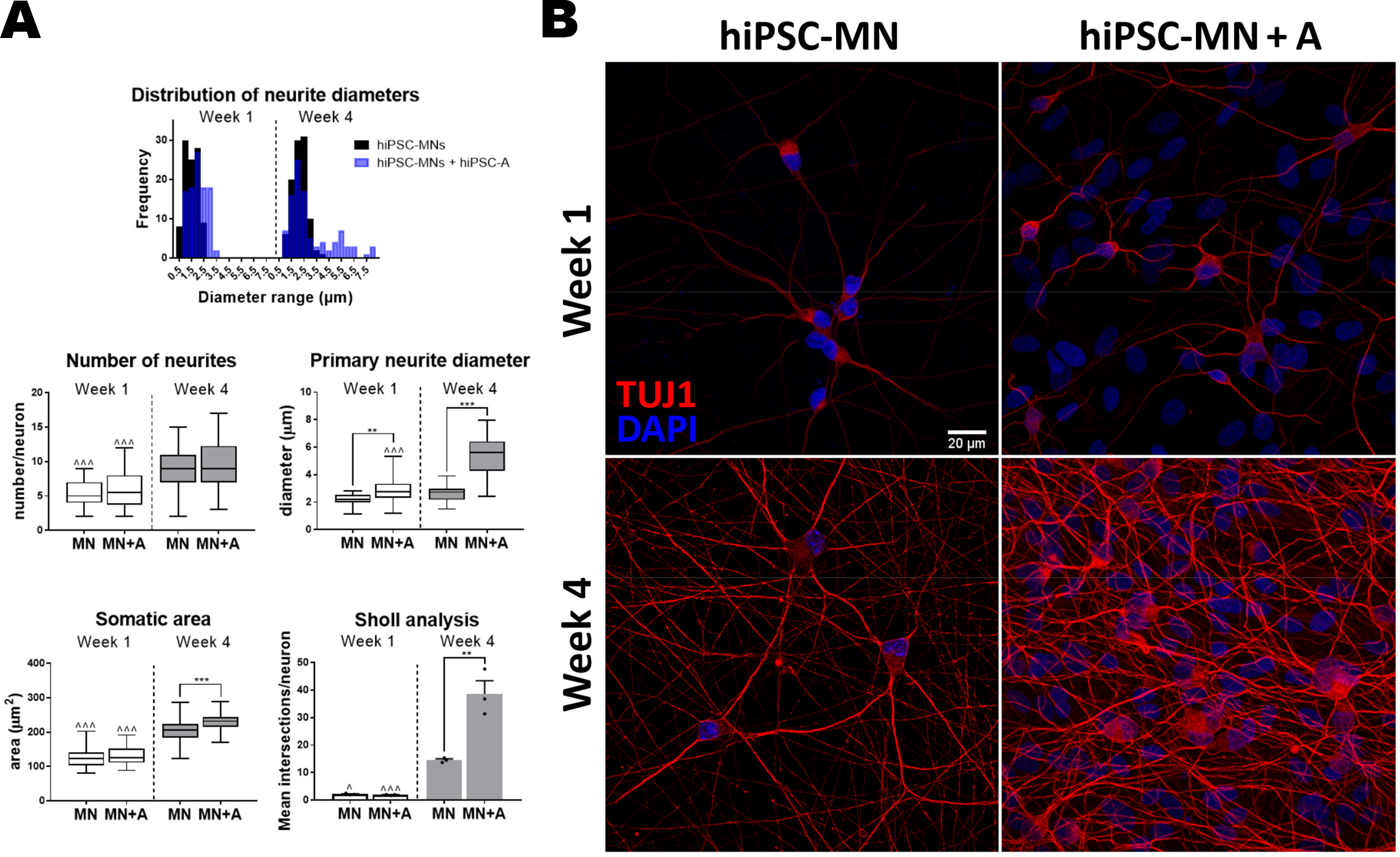
Morphological analysis of hiPSC-MN maturation with hiPSC-A co-culture. **A**: Quantification of neuronal morphology between hiPSC-MN in mono-culture (MN) or in co-culture with hiPSC-A (MN+A). Distribution (histogram) refers to the diameters of *n*=100 neurites per condition at 1 and 4 weeks in culture. Number of neurites, primary neurite (i.e. largest neuronal process) diameter and somatic area were determined analyzing *n*=50 neurons per culture condition and time-point. Sholl analyses results are shown as mean number of intersections per neuron for *n*=3 coverslips per condition, with a minimum of 10 neurons analyzed per coverslip. Experimental conditions (mono-vs co-cultures) are compared within each time-point (*), i.e. week 1 (white bar) or week 4 (grey bar) after plating; similarly, time-point observations (week 1 vs week 4) are compared within each condition (^ on the top of week 1 bars). * *or* ^ *p*<*0.05;* ** *or* ^^ *p*<*0.01;* *** *or* ^^^ *p*<*0.001*. **B**: Representative immunofluorescence images utilized for morphological analyses of hiPSC-MN, in mono or co-culture, at week 1 and 4 after plating.

Through parallel qPCR experiments we sought to verify our observations from immunostaining and to examine the effect of human astrocyte maturity as well as species-specific interactions on neuronal maturation (Figure 5). We confirmed that spinal cord motor neuron identity, as defined by ChAT expression, is not significantly influenced by hiPSC-A co-culture at either 60 or 90 DIV nor is it influenced by co-culture with mouse astrocytes. In contrast, the expression of genes required for the biosynthesis of neurotransmitter receptors (GRIA2, 3 for AMPA receptor, GABRA1 for GABA receptor, and GLRA1 for glycine receptor) and of GABA (GAD1) is particularly enriched in the co-cultures with hiPSC-A, in a time-dependent manner (Figure 5). We found that even though the presence of immature or mouse astrocytes increased neuronal maturation when compared to hiPSC-MN monocultures, particularly at the later time point, this effect was consistently lower than that observed for co-cultures with mature (i.e. cultured for 90 DIV prior to use) and human-derived astrocytes (Figure 5).

**Figure 5.**
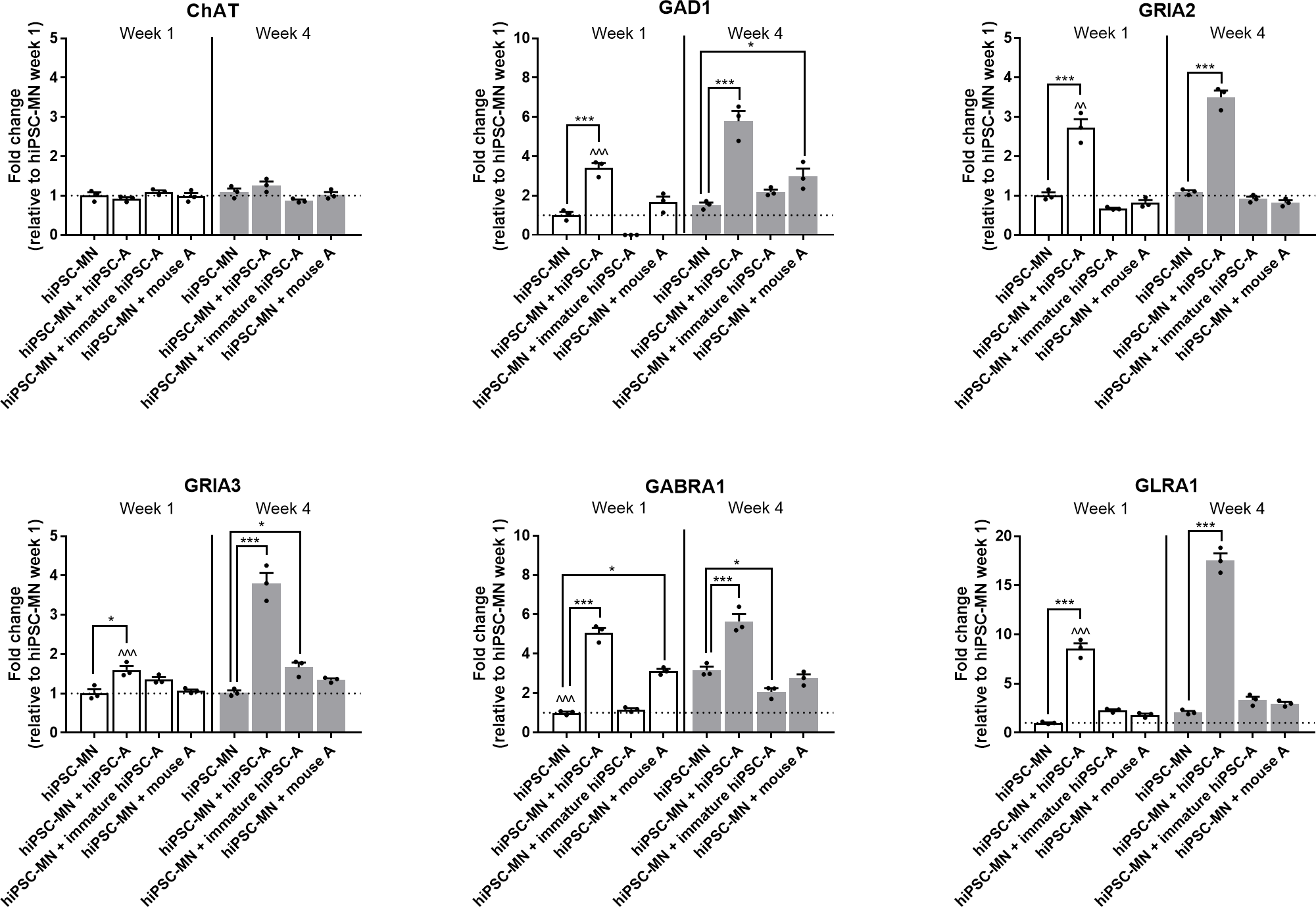
qPCR analysis of neuronal RNA transcripts. Markers of neuronal subtypes (ChAT, GAD1) and neurotransmitter receptors subunits (GRIA2, GRIA3, GABRA1, GLRA1) (mean of *n*=3 samples per condition). Experimental conditions, i.e. hiPSC-MN mono-cultures *vs* co-cultures of hiPSC-MN with mature hiPSC-A (“hiPSC-A”), with “immature hiPSC-A” or with mouse primary spinal cord astrocyte (“mouse A”), are compared within each time-point (*). Time-point observations (week 1 vs week 4) are compared within each condition (^). * *or* ^ *p*<*0.05;* ** *or* ^^ *p*<*0.01;* *** *or* ^^^ *p*<*0.001*.

### hiPSC-MN influence hiPSC-A maturation

We sought to investigate astrocyte-neuron interactions in our human spinal cord iPSC-based platform, as their reciprocal maturation could dynamically influence the electrophysiological activity of hiPSC-MN recorded by MEA.

Confirming previous descriptions by Roybon and colleagues^14^ and by our group^15^, we defined a maturing glial identity of our hiPSC-A, as shown by immunoreactivity of cells for S100β (78.9±4.2%), GFAP (41.2±3.3), EAAT2 (8.1±1.6%) and Cx43 (22.7±2.5%) (Figure 6). Spinal cord regional identity was suggested by high immunoreactivity (92.5±1.4%) for HOXB4 (Supplementary figure 1B). These astrocytes did not show relevant immunoreactivity for CD51 (4.1±0.2%), a marker recently associated with cortical identity ^33^ (Supplementary figure 1B).

**Figure 6.**
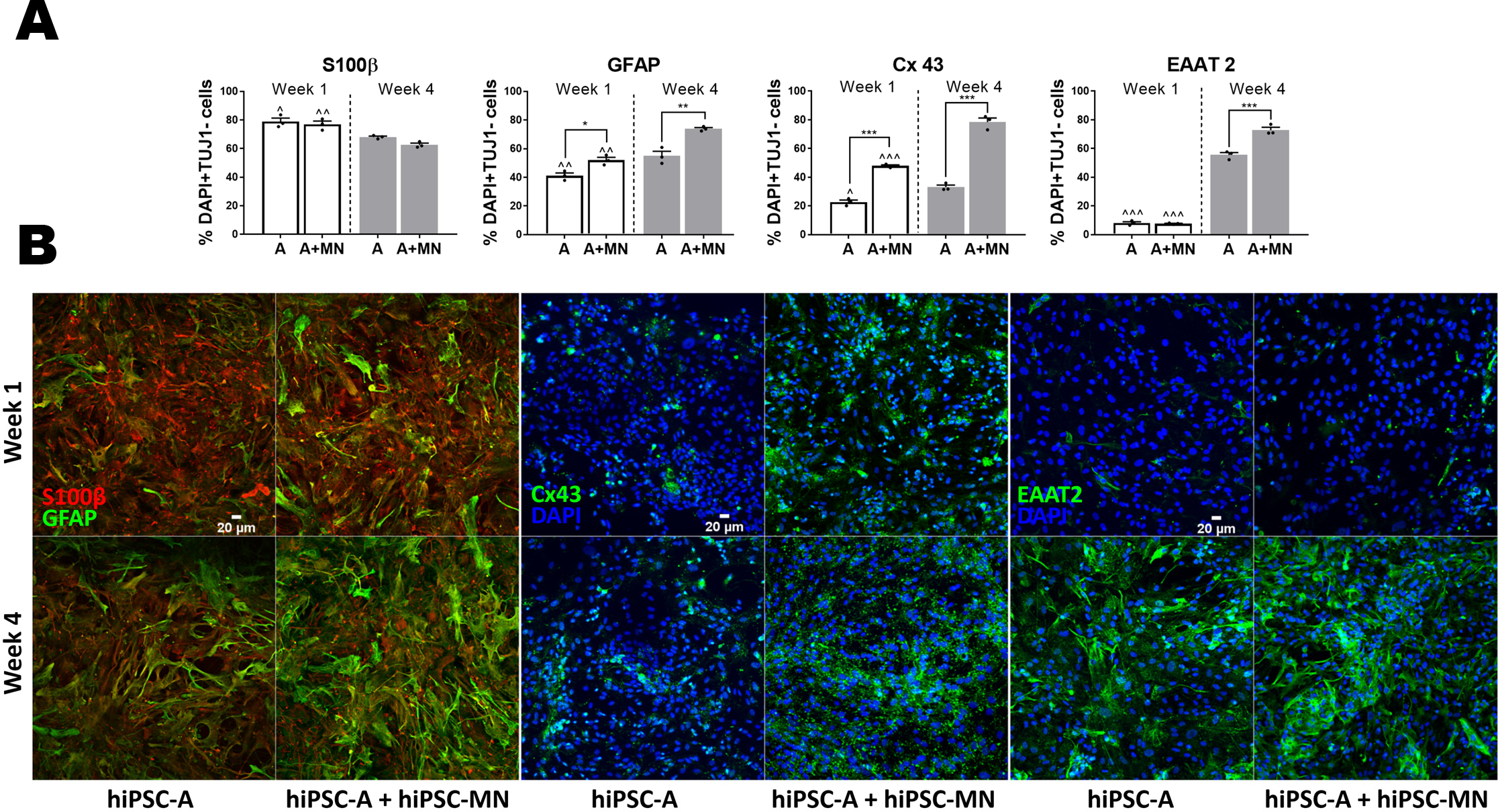
**A: Immunofluorescence quantification of astrocyte-specific markers.** We considered the total number of DAPI^+^ cells which were negative for TUJ1 immunostaining as an approximation of the number of astrocytes in hiPSC-A+MN co-cultures (A+MN), and hiPSC-A mono-cultures (A); astrocyte-specific markers are expressed as percentage of DAPI^+^ and TUJ1^−^ cells. S100β^+^ marks a less mature stage of astrocytic differentiation. Conversely, GFAP, Cx43 and EAAT2 are markers of maturing/mature astrocytes. Experimental conditions (hiPSC-A mono-cultures vs hiPSC-MN + hiPSC-A co-cultures) are compared within each time-point (*). Time-point observations (week 1 vs week 4) are compared within each condition (^). * *or* ^ *p*<*0.05;* ** *or* ^^ *p*<*0.01;* *** *or* ^^^ *p*<*0.001)*. **B**: **Representative immunohistochemical images of astrocyte markers.** Reciprocal changes of S100β and GFAP according to culture conditions and time in vitro. Immunoreactivity for Cx43 and EAAT2 in hiPSC-A mono- and co-culture, at week 1 and week 4 after plating.

We then found that the co-culture of hiPSC-A with hiPSC-MN, when compared with hiPSC-A mono cultures, increased the immunoreactivity for GFAP (52.0±3.5%, p<0.01), and Cx43 (47.9%±1.2, *p*<0.001) (Figure 6), thus suggesting a more mature astrocyte phenotype. This enhanced glial maturation was even more marked after 4 weeks of co-culture, when, in addition to GFAP (73.9±1.6% vs 54.9±5.6, *p*=0.001) and Cx43 (78.5±4.8% vs 47.9±1.2%, *p*<0.001), EAAT2 immunoreactivity increased significantly (72.9±3.5% vs 55.6±2.9%, p<0.001) (Figure 6).

QPCR experiments confirmed the immunostaining findings at mRNA expression levels on a larger set of astrocyte genes, which included also AQP4 and ALDH1L1 (Figure 7). We used this platform to examine less mature hiPSC-A cultured for only 60 DIV prior to use, and to investigate their maturation in the presence of hiPSC-MN. One week after their plating in co-culture, qPCR experiments suggested an immature status of this glial population when compared to “mature” (i.e. 90DIV) hiPSC-A in mono and co-culture (Figure 7). After 4 weeks of co-culture, these cells remained significantly less mature, particularly when considering EAAT2 and GFAP expression levels, which were negligible. AQP4 and particularly Cx43 were two exceptions, as their expression levels in immature hiPSC-A were comparable for AQP4, or even higher for Cx43 (*p*<0.001), than the more mature hiPSC-A (cultured for 90DIV prior to use) in mono or co-culture (Figure 7).

**Figure 7.**
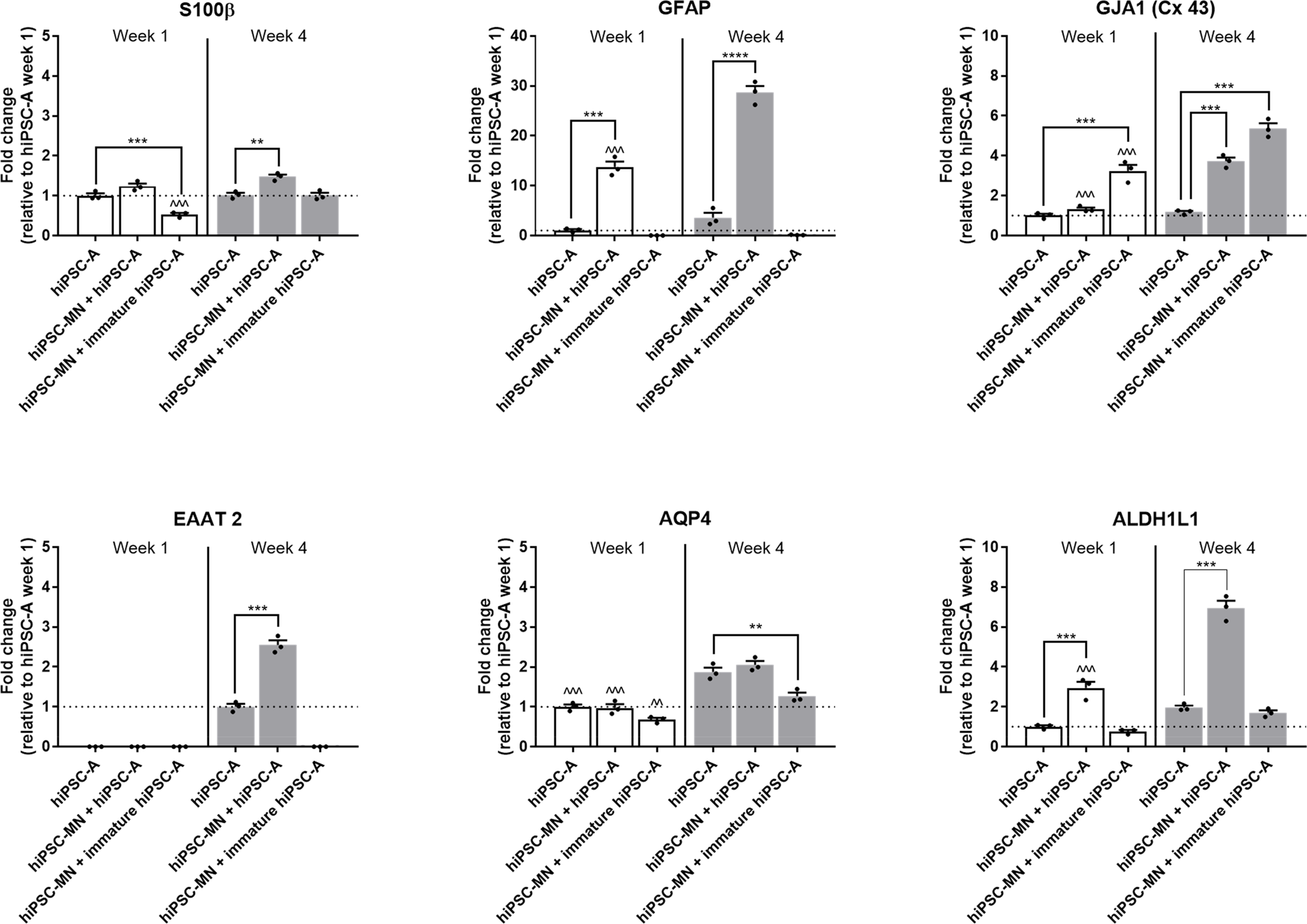
qPCR analysis of astrocyte RNA transcripts. Expression of immature (S100β) and more mature (GFAP, GJA1, EAAT2, AQP4 and ALDH1L1) astrocyte transcripts were quantified (mean of *n*=3 samples per condition). Experimental conditions (hiPSC-A mono-cultures *vs* co-cultures of hiPSC-MN with mature hiPSC-A or with immature hiPSC-A) are compared within each time-point after plating (*). Time-point observations (week 1 vs week 4) are compared within each condition (^). * *or* ^ *p*<*0.05;* ** *or* ^^ *p*<*0.01;* *** *or* ^^^ *p*<*0.001*.

### MEA models hiPSC-A and hiPSC-MN interactions

Given the striking hiPSC-A dependent changes in hiPSC-MN maturation by immunofluorescence and qPCR, we sought to determine whether these structural changes and altered expression profiles are meaningfully related to altered electrophysiologic function of neurons. We plated hiPSC-MN in monocultures or co-cultures with astrocytes on MEA plates that were either plated concurrently or serially. We recorded from the MEA plates at 1, 2, 3, and 4 weeks after plating (Figure 8), corresponding to 67, 74, 81, 88 DIV and 97, 104, 111, 118 DIV for hiPSC-MN and hiPSC-A, respectively. We noted increases in all electrophysiological parameters evaluated (spike rate, burst rate, percentage of electrodes spiking and percentage of electrodes bursting) over time (Figure 8A-D). The changes over time occurred for all culture conditions (hiPSC-A/hiPSC-MN simultaneous co-culture, hiPSC-MN followed by hiPSC-A, hiPSC-A followed by hiPSC-MN, and hiPSC-MN culture alone) (*p*<0.001 for week 1 vs. week 4 comparisons). However, the difference in the maturation between simultaneously cultured hiPSC-A and hiPSC-MN when compared with hiPSC-MN cultured alone was most dramatic with the spike rate, burst rate, number of spiking and number of bursting electrodes higher in the simultaneous co-culture at all time-points. Though less marked and uniform over time-points, the simultaneous co-cultures also showed greater degrees of neuronal activity when compared to serial co-cultures (Figure 8A-D). We did not see any evidence of spontaneous synchronous bursting activity which is thought to reflect high degrees of neuronal connectivity^17,21^ (not shown). For all conditions, MEA plates with hiPSC-A alone were recorded as controls, and did not show any measurable electrophysiological activity within the 4-week period of recording, confirming the relatively pure composition of our astrocyte cultures (Supplementary Figure 1B). We calculated the percentage of the 59 active electrodes whose spike activity persisted throughout the 4 weeks of recordings, and defined this parameter as “consistently active electrodes” (Figure 8E). We noted that electrophysiological consistency was influenced by the sequence of culturing hiPSC-A and hiPSC-MN. Indeed, approximately 45.8±4.4% of electrodes remained persistently active over 4 weeks following the simultaneous culture of iPSC-A and iPSC-MN. The culture of either hiPSC-A first (25.4±3.4%) or hiPSC-MN first (27.1±4.5%) were not significantly different from each other, but were lower than the simultaneous culture (*p*<0.05). Furthermore, only 13.0±4.2% of electrodes showed persistent spike activity when hiPSC-MN were cultured in the absence of hiPSC-A (Figure 8E) (*p*<0.001).

**Figure 8.**
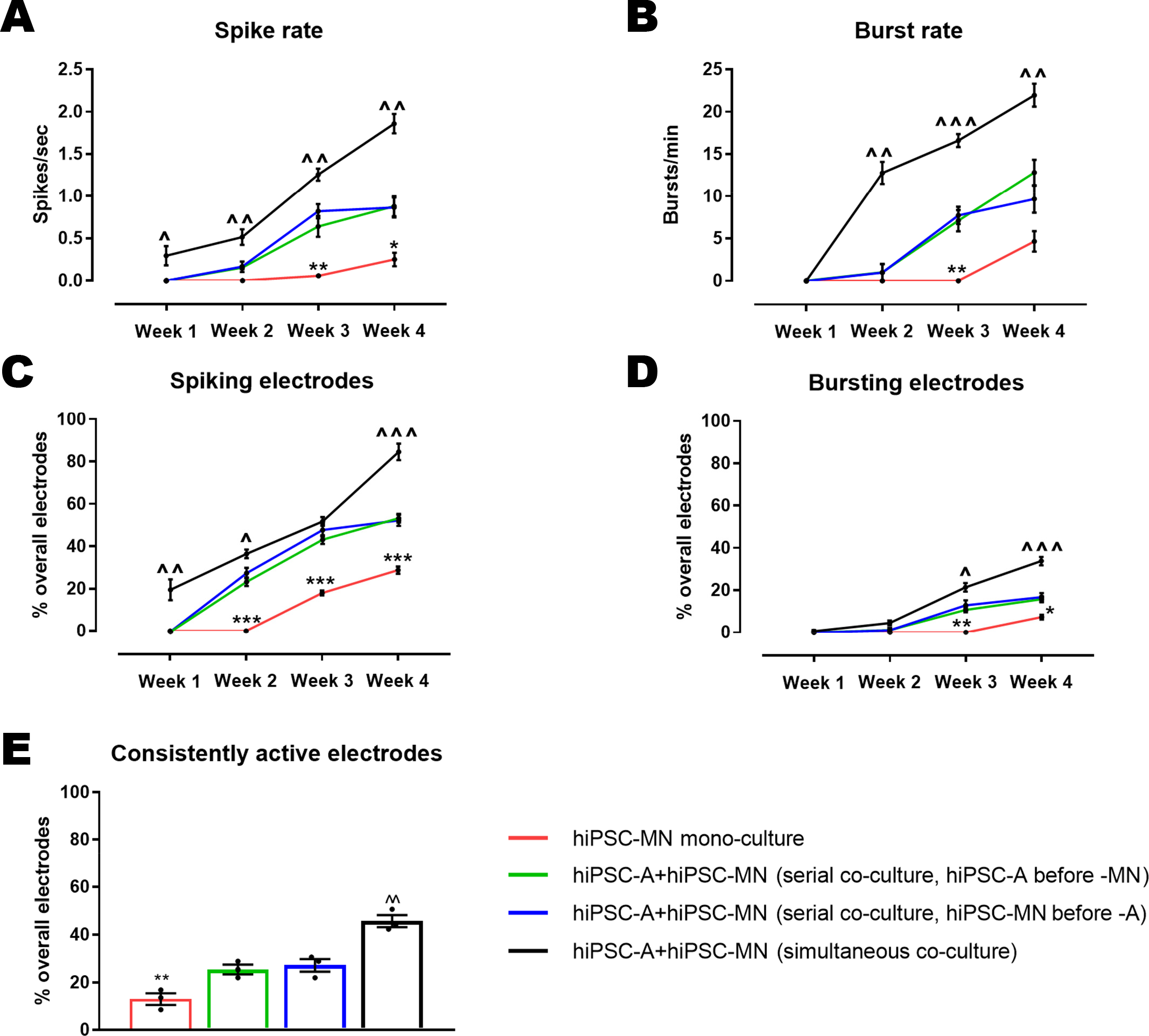
The electrophysiological maturation of hiPSC-MN by hiPSC-A over time. **A-D**: Electrophysiological parameters were recorded at weekly intervals for hiPSC-MN in mono-culture as well in co-culture with hiPSC-A. Data are presented as mean±SEM (mean of *n*=3 MEA plates). **E**: The percentage of consistently active electrodes represents populations of neurons with stable electrophysiological activity over individual electrodes (*n*=3 cultures). Comparisons of hiPSC-MN monocultures with the two hiPSC-MN/A serial co-cultures (*). Comparisons between the simultaneous co-culture and the two serial co-cultures (^). * *or* ^ *p*<*0.05;* ** *or* ^^ *p*<*0.01;* *** *or* ^^^ *p*<*0.001*.

In addition to the 4 week recordings, we tested the long-term survival of our optimized co-culture, and succeeded in maintaining these cells in culture for up to 9 months. A trending increase in spike rate was noted through the first 5 months which then plateaued (Supplementary Figure 2). This is consistent with data derived from cortical hiPSC-neurons^7^. The feasibility of this long-term culture system is instrumental to future studies investigating the electrophysiological, morphological and molecular properties of hiPSC-A/-MN co-cultures at stages of their maturation that more closely resemble their *in vivo* counterparts.

We also examined whether the plating density of hiPSC-A influences hiPSC-MN activity (Figure 9A). The spike rate, percentages of electrodes exhibiting spikes and percentages of electrodes exhibiting bursts were significantly increased across all time-points in co-cultures with a density of 1×10^5^ astrocytes/plate when compared with a lower density of 0.5×10^5^ astrocytes/plate. However, when even higher densities of astrocytes (2×10^5^ astrocytes/plate) were utilized, there was poor cell adhesion immediately after plating, which led to lifting of the cultures (not shown). These findings were correlated with a higher percentage of “consistently active electrodes” in the co-cultures with 1×10^5^ astrocytes/plate when compared with a lower density of 0.5×10^5^ astrocytes/plate (43.6±2.4% vs 28.8±2.6%, *p*<0.05).

**Figure 9.**
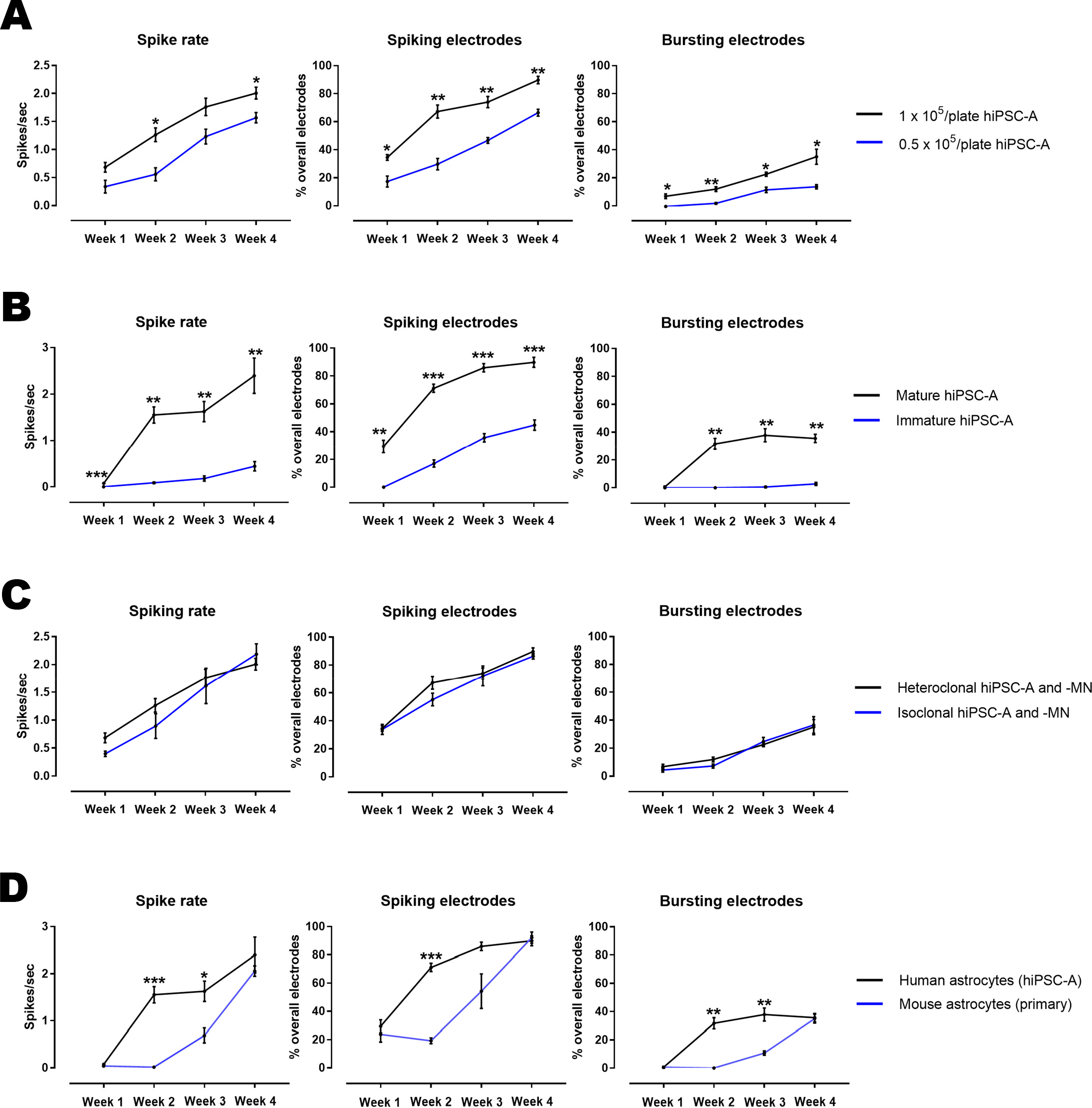
Astrocyte variables influencing the electrophysiological maturation of hIPSC-MN in co-culture. **A**: Influence of hiPSC-A density on the electrophysiological maturation of hiPSC-MN as measured by MEA. **B**: Effect of immature vs mature astrocytes on neuronal maturation as measured by MEA. **C**: Influence of hiPSC-A and hiPSC-MN respective clonal origin on the electrophysiological maturation of hiPSC-MN. **D**: Effect of the species origin (mouse primary spinal cord astrocytes vs human iPSC-A) on hiPSC-MN maturation, as recorded by MEA. Data are the mean of *n*=3 MEA plates for all conditions. \**p*<*0.05,* \*\**p*<*0.01,* \*\*\**p*<*0.001*.

To assess whether astrocyte age and maturity prior to co-culture with hiPSC-MN would influence hiPSC-MN electrophysiological activity, we cultured hiPSC-A for 60 days prior to the co-culture with hiPSC-MN. When compared to co-cultures with mature (90DIV) hiPSC-A, these cells had a significantly reduced capacity to support hiPSC-MN spike rate and number of active electrodes (Figure 9B). This phenomenon persisted over the 4-week time course (*p*<0.05 for all comparisons).

Given that using hiPSC raises the question as to whether there could be variations in electrophysiological properties amongst different iPSC lines, we differentiated astrocytes and MN from a single normal individual (GM01582) (“isoclonal” hiPSC-A and -MN). We then differentiated astrocytes from another normal individual (CIPS) and co-cultured those hiPSC-A with hiPSC-MN (GM01582) (“heteroclonal” hiPSC-A and –MN). After recording weekly for the 4 weeks of co-culture, we did not appreciate significant clone-specific differences in electrophysiological activity (Figure 9C).

Most studies utilizing MEA for recording from human iPSC-neurons have employed rodent astrocytes in the context of co-culture systems^7,9^. We wanted to determine whether rodent astrocytes would alter the electrophysiological maturation of hiPSC-MN. Our results demonstrate that mouse spinal cord astrocytes result in a slower maturation of the electrophysiological activity of hiPSC-MN when compared to co-cultures with hiPSC-A. This is most notable at the earliest recorded time points (Figure 9D).

### MEA as a screening platform for drugs targeting hiPSC-MN and hiPSC-A

Beyond the spontaneous electrophysiological activities of hiPSC-MN, we examined the response of these cells, in mono- or co-culture to compounds targeting ion channels and neurotransmitter receptors. When cultured alone, hiPSC-MN responded to the application of a non-specific depolarizing agent, such as 100 mM of potassium chloride (KCl), with an increase in spike rate without any change in the number of spiking or bursting electrodes (Supplementary Figure 3). However, the application of neurotransmitter agonists/antagonists at appropriate working concentrations, including KA (5μM), CNQX (50μM), and bicuculline (10 μM) did not alter MEA electrophysiological parameters (Supplementary Figure 3).

In contrast, when we tested these compounds on hiPSC-A/hiPSC-MN simultaneous co-cultures (Figure 10), we found that the AMPA/kainate receptor agonist, KA, and antagonist, CNQX, induced appropriate (i.e. an increase for KA and a decrease for CNQX) changes in the spike rate together with a significant variation of bursting activity; these responses were time-dependent, as they were absent at earlier stages of maturation in vitro (week 1-2 after plating), while they appeared after 3-4 weeks of co-culture. The addition of 10μM concentrations of the GABA antagonist, bicuculline, did not significantly change the overall MEA activity, though there was a trend towards increased spike rate and percentage of bursting electrodes in older co-cultures (Figure 10). In contrast to the neurotransmitter modulators, KCl showed depolarizing effects across all time points (Figure 10). We did not observe any hypersynchronous bursting activity occurring after the application of any of these compounds.

**Figure 10.**
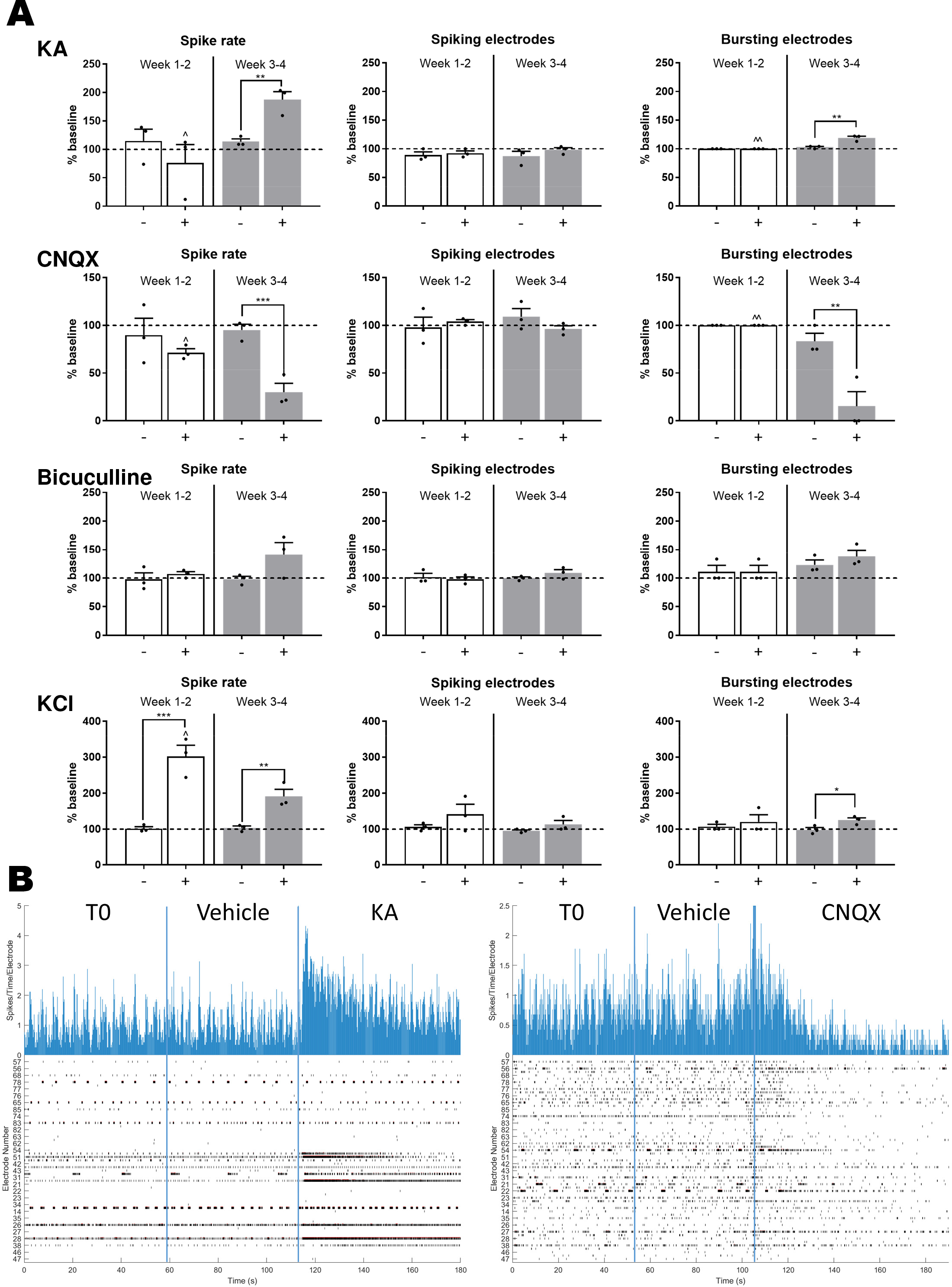
hiPSC-A influence the responses of hiPSC-MN to neurotransmitters. **A**: hiPSC-MN responses to compounds acting on neurotransmitter receptors (KA, CNQX, and bicuculline) and to a non-specific depolarizing agent (KCl), according to culture condition (mono-vs co-culture with hiPSC-A) and over time (week 1-2 after plating - white bars, and 3-4 after plating - grey bars). The activity recorded by MEA plates after vehicle addition (−) and after drug addition (+) was normalized to the baseline activity (horizontal dashed line) (mean of n=3 independent experiments per drug and time point). Comparisons between activity after vehicle and after drug addition (*). Time-point observations (week 1-2 vs. week 3-4) are compared within each condition (^). * *or* ^ *p*<*0.05;* ** *or* ^^ *p*<*0.01;* *** *or* ^^^ *p*<*0.001*. **B**: Sample of real-time recording of hiPS-MN/hiPSC-A co-cultures by MEA. Two representative tracings of electrophysiological activity are shown: the effect of the addition of KA (*left*) and CNQX (*right*) compared to the addition of the vehicle and to baseline activity. Top histograms represent the spike rate per electrode and per time point of recording, while the bottom traces represent overall electrodes showing spiking activity per time point. Abbreviations: KA, kainic acid; KCl, potassium chloride.

To examine the immediate effect of altered astrocyte function on neuronal firing, we used our MEA platform to test compounds targeting astrocyte specific channels, and evaluate their indirect influence on neuronal firing. The application of 340μM GAP19, a connexin 43 hemichannel blocker, resulted in a significant reduction of MEA activity (spike rate and bursting electrodes). This phenomenon was apparent across all time points of co-culture and was dose-dependent (Figure 11). To ensure that this effect was mediated by astrocytes, we tested the drug on hiPSC-MN mono-cultures, and did not appreciate any variation in neuronal activity (Supplementary Figure 3). The selective EAAT2 glutamate transporter antagonist, DHK, caused a reduction in MEA activity, which appeared within 10 minutes of recording. This effect was dose dependent, and could be appreciated only in co-culture at later time points (week 3-4) (Figure 11).

**Figure 11.**
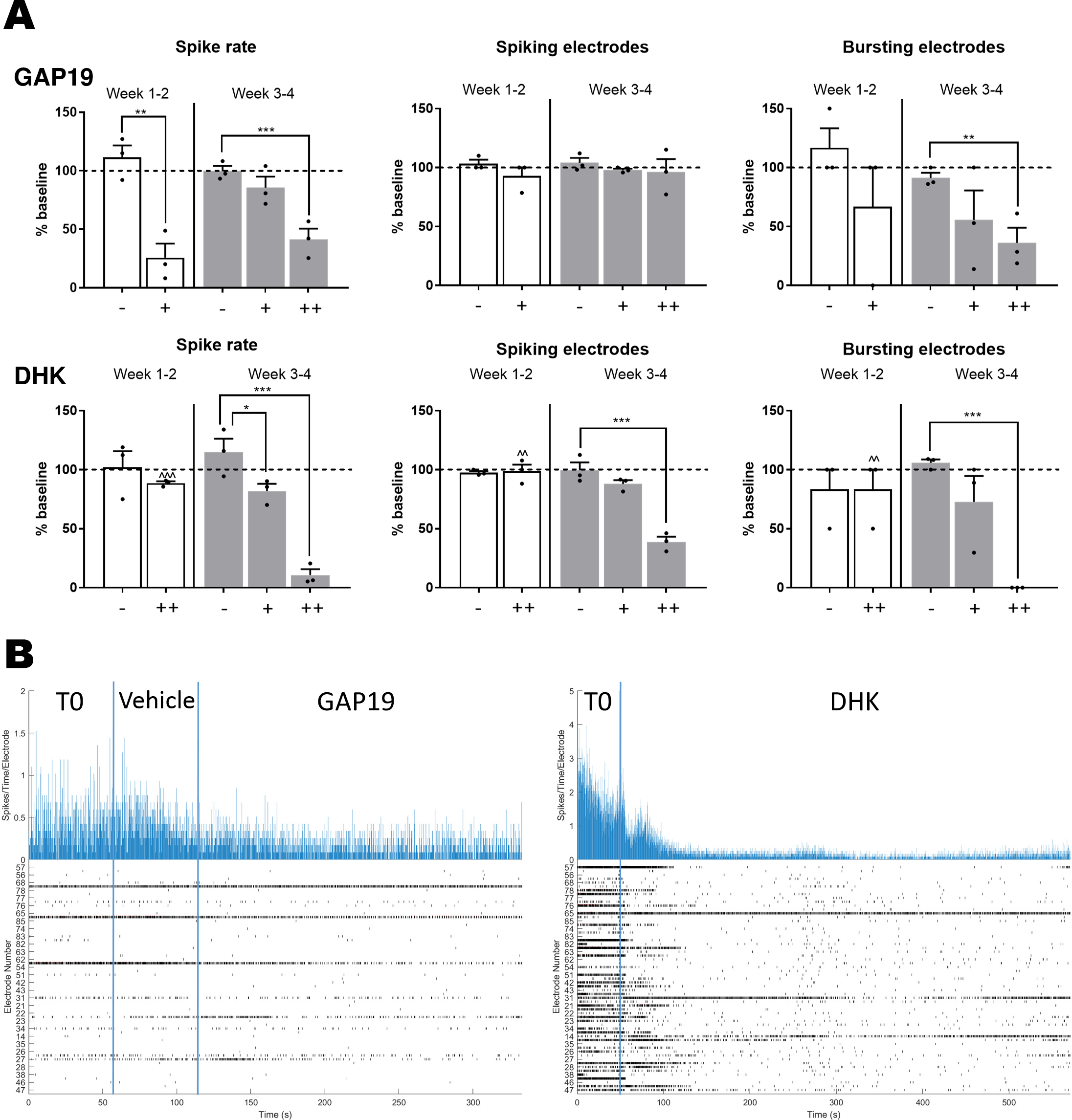
Compounds targeting astrocytes affect hiPSC-MN electrophysiology as recorded by MEA. **A**: Effects of GAP19, a specific Cx43 hemichannel blocker, on neuronal activity, according to culture condition (mono-vs co-culture with hiPSC-A) and over time (week 1-2 after plating - white bars, and 3-4 after plating - grey bars) (- vehicle, + 34 μM, ++ 340 μM). Changes on hiPSC-MN electrophysiological activity after the addition of DHK (- vehicle, + 50μM, ++ 300μM). The electrophysiological parameters were normalized to the baseline activity recorded for 1 minute (dashed line) (mean of *n=3* independent experiments per drug and per time point*) (*\* *or* ^ *p*<*0.05;* ** *or* ^^ *p*<*0.01;* *** *or* ^^^ *p*<*0.001)*. **B**: Representative MEA tracings of electrophysiological activity following the addition of GAP19 (left) and DHK (right). Abbreviations: DHK, dihydrokainic acid

## Discussion

To our knowledge, our current study is the first to investigate how hiPSC-derived glia, differentiated into a distinct spinal cord astrocyte identity^14,34^, influence the morphological, molecular, electrophysiological and pharmacological properties of hiPSC-MN in a MEA-based platform. The regional specificity of this paradigm is particularly relevant as advances in the generation of cell subtypes from human iPSCs are being used in precision medicine strategies for modeling neurological diseases. Previous literature on MEA applied to hiPSC studies have relied exclusively upon hiPSC-derived cortical neurons, for *in vitro* modeling of epilepsy and seizures^8,9^. Though recent MEA studies have utilized neurons co-cultured with human iPSC-derived astrocytes^5,8–10^, less attention has been given to how astrocytes may influence neuronal electrophysiological activity—particularly in the context of human biology. Bidirectional interactions between hiPSC-MN and hiPSC-A may be hypothesized in this MEA platform, but they have not yet been investigated.

To establish a hiPSC platform of spinal cord astrocyte/MN co-culture, we first optimized the sequence of co-culture. As observed by others^35^, the culture of hiPSC-MN in the absence of astrocytes resulted in a significant number of large aggregates of iPSC-MN that morphologically made it difficult to discern specific boundaries and connections amongst these cells. Furthermore, these cultures demonstrated diffuse neuronal mobility, as detected electrophysiologically by significant variability in recordings from individual MEA electrodes. This is reflected in the observation that only 13% of electrodes had activity which was persistent for the 4-week recording period. The culture of either hiPSC-A followed by plating of hiPSC-MN or the converse, resulted in fewer neuronal aggregates and an increase in the number of MEA electrodes showing persistent activity. Finally, we observed that the simultaneous plating of hiPSC-A and hiPSC-MN morphologically resulted in a much more homogeneous distribution of hiPSC-MN, and in the greatest number of electrodes showing persistent activity.

Important to utilizing this human spinal cord-specific co-culture system was ensuring that hiPSC-derived neuron identity did not change with time in culture, nor did the density of either astrocytes or neurons, as this could influence the interpretation of MEA recordings. Our data suggest that the majority of our cells are spinal cord motor neurons (positive for ChAT and, to a lesser extent, for ISL1/2), expressing appropriate neurotransmitter receptors, with AMPA receptors being more represented than GABA and glycine receptors, as would be expected in the spinal cord. Importantly, in accordance with our protocol of ventralization and caudalization, and in contrast to previous studies involving hiPSC-cortical neurons^8,9^, only a very small percentage of neurons were corticospinal CTIP2^+^ motor neurons.

The observation that neuronal subtypes and cell densities (both glial and neuronal) did not change over time, made us more confident that the neuronal activity of these cells, as recorded by MEA, was not a reflection of changing neuronal populations but rather a representation of neuronal maturation. The stable composition of hiPSC-derived neuronal subtypes over time has been observed by others as well ^32^, and may reflect an early determination of the neuronal fate during NPCs differentiation.

In accordance with previous studies examining hiPSC-cortical neurons^7,9^, hiPSC-MN maturation over-time was mirrored by morphological changes, with increases in neurite number, size, and complexity accompanied by an increase in the number of synapses. These changes were enhanced by spinal cord hiPSC-A co-cultured with MN.

We used this optimized co-culture platform for MEA recording in order to examine hiPSC-A contributions to motor neuron physiology. We first observed that astrocytes influenced hiPSC-MN electrophysiology over time with increases in firing rate and burst frequency that far exceeded the activity of hiPSC-MN cultured in the absence of astrocytes. These findings are consistent with previous observations in other cell neuronal subtypes suggesting that astrocytes promote electrophysiological maturation of hiPSC-neurons^5,10,36–42^.

We then determined that the ratio of hiPSC-A to hiPSC-MN influenced the activity of MN with higher concentrations of hiPSC-A providing a significant increase in hiPSC-MN firing. We have previously described that hiPSC-A require longer culture times *in vitro* in order to acquire more mature astrocyte markers^15^. The significantly lower hiPSC-MN firing when cultured with 60-day astrocyte cultures, which was paralleled by qPCR profiling of hiPSC-MN, suggests that for the purposes of examining electrophysiological activity, more mature astrocytes seem to be essential.

One of the concerns regarding the use of human iPSCs, as opposed to rodent astrocytes, for MEA studies is that genetic heterogeneity amongst individuals is much greater than in rodents thus making interpretation of results more challenging. We utilized control hiPSC-A lines from 2 unrelated individuals and cultured them with a single hiPSC-MN line to address whether the clonal origin of astrocytes and neurons influenced hiPSC-MN electrophysiological activity. We did not appreciate differences in the electrophysiological variables over time between the 2 control hiPSC-A lines which is in contrast to a previous study which found that the clonal origin of hiPSC-A influenced neuronal activity^43^. While there are a multitude of potential combinations of hiPSC-A lines that could be studied, our data provide early evidence that the source of control iPSC-derived astrocytes does not significantly influence iPSC-MN activity.

Given that hiPSC-A differentiation requires longer time in culture, generates less mature astrocytes, and produces fewer astrocytes than primary rodent-derived astrocytes, we examined whether there was a difference in neuronal activity of co-cultures with hiPSC-A compared to mouse primary spinal cord astrocytes. We found that rodent spinal cord astrocytes resulted in a delayed pattern of hiPSC-MN electrophysiological maturation when compared to hiPSC-A. This difference paralleled qPCR profiling showing that the maturation of hiPSC-MN was more robust when human astrocytes were utilized. This appears to be related to species differences rather than regional heterogeneity since rodent astrocytes were derived from the spinal cord. Our findings are in contrast to those of Lischka and colleagues^43^ who found that neonatal mouse cortical but not isogenic human astrocyte feeder layers enhanced the maturation of iPSC-N. It must be noted, however, that those investigators used a cortical patterning protocol to differentiate iPSC-N, mouse forebrains to derive rodent astrocytes, and a patch-clamp platform to record neuronal activity. Our observations may not be surprising as a growing body of evidence suggests that human astrocytes are more complex than their rodent counterparts, both morphologically ^44^ and physiologically ^44,45^. Beyond stressing the importance of standardizing cell culture conditions and electrophysiological recording platforms, our data suggest that human and regional identity of astrocytes may be relevant factors that influence their cross-talk with neurons.

Essential to the validation of this co-culture strategy was the demonstration that hiPSC-MN electrophysiological activity was not merely spontaneous but would also respond to the application of neurotransmitters and that their activity could be inhibited by receptor antagonists. Consistent with the identity of these cells as iPSC-MN, in our co-culture platform, we observed a robust and consistent response to the glutamate agonist, kainic acid, and the AMPA antagonist, CNQX. As shown by others for hiPSC-cortical neurons^9,10^, this effect was maturation-dependent, as it appeared only after 3-4 weeks *in vitro*, while it was absent at earlier stages of maturation. The response to the GABA antagonist bicuculline was less robust, a finding consistent with a lower representation of GABA receptors. In contrast, neuronal cultures without astrocytes did not show any response to AMPA nor to GABA agonists/antagonists, which is consistent with our observations on the immaturity of hiPSC-MN monocultures, and parallels previous findings on hiPSC-cortical neurons^5,9,35^.

The few previously published studies on MEA recording from co-cultures of hiPSC-A/hiPSC-N have been carried out using human iPSC-derived cortical neurons^5,8,9^. Consistent with our observations, these studies have demonstrated that human astrocytes enhance different MEA parameters including: spiking rate, bursting rate and number of spiking and bursting electrodes. However, these studies utilizing cortical neurons recording have demonstrated some electrophysiological patterns, such as synchronized burst firings^5,9^, which are indicative of high connectivity and synchronization among neurons and may model the network activity of the cortex in its normal functioning or in pathological states such as epilepsy^21^. We did not appreciate similar degrees of connectivity in hiPSC-MN firing up to 9 months of culture *in vitro*. Given that morphologically we see evidence of neurite outgrowth generating complex networks and connecting neurons through well-developed synapses, the lack of this form of connectivity may suggest that the spinal cord neuronal lineages are not capable of a similar synchronous activity and, thus, truly model spinal cord neural population.

Interestingly, the morphological and molecular characterization of our co-culture platform showed that astrocyte maturation, as defined by different glial-specific markers, was enhanced by the presence of neurons. We also found that these effects were time dependent and marker dependent, as shown by the profile of maturation of “immature” hiPSC-A (i.e. hiPSC-A cultured for 60DIV prior to use) in co-culture, which was less marked than 90DIV hiPSC-A, with, however, the exception of two markers, Cx43 and AQP4 which appeared to be influenced by the presence of neurons at a very early time points of astrocyte maturation. These early changes in AQP4 and Cx43 profiles suggest a neuronal influence on their expression, not related to astrocyte maturation. In this respect, and beyond our preliminary observations based on RNA levels, previous literature on rodent astrocytes had already shown that Cx43 and AQP4 function is modulated by neuron-astrocyte interactions, as suggested by elevations in calcium intercellular signaling^46,47^ and changes in astrocyte volume^48,49^, respectively, in response to neuronal activity.

To our knowledge, spinal cord astrocyte influences on motor neuron maturation^50^ have not yet been investigated in a fully human iPSC co-culture. In a previous study^15^, we analyzed the transcriptomic profile of hiPSC-A after in vivo transplantation in the rat spinal cord, and found that of the top 20 astrocyte-specific genes reported in the literature, approximately half of these were increased after transplant, with GFAP, EAAT2, CX43 and AQP4 most abundantly expressed^15^. Besides confirming these previous findings in a fully human hiPSC-based *in vitro* platform, our data suggest that iPSC-A maturation in the spinal cord microenvironment may be, at least partially, determined by the presence of neurons.

Astrocyte-neuron cross-talk is relevant to our MEA platform since hiPSC-MN electrophysiological maturation may be, at least partially, influenced by the concurrent astrocyte maturation in co-culture. In this regard, the expression profile of immature hiPSC-A in co-culture with hiPSC-MN was paralleled by lower degrees of neuronal activity on MEA. Furthermore, the pharmacological effects of GAP19 and DHK on MEA activity paralleled Cx43 and EAAT2 expression, respectively, by hiPSC-A in co-culture, with GAP19 being active in both early and late co-cultures (both characterized by high expression of Cx43), and DHK showing significant effects only after 3-4 weeks of co-culture (when EAAT2 expression became relevant). Although it is well known that neuronal firing exerts multiple effects on astrocytic gap-junctions^51^, we show that a converse influence may occur, since the pharmacological manipulation of Cx43 by GAP19, a hemichannel blocker, resulted in marked changes in neuronal activity. This phenomenon was astrocyte-specific, as GAP19 did not influence hiPSC-MN firing in cultures without astrocytes. The inhibitory effect of DHK on neuronal firing, and particularly on burst activity, was unexpected, but confirms previous findings on rodent cell cultures and rodent brain slice preparations^21,22^. This may be the result of an alteration of the recycling of glutamate for pre-synaptic release^21^, or of the inhibitory action of the pre-synaptic metabotropic glutamate receptor (mGluR)^22^. Although further experiments may be needed to clarify the mechanism of these glial influences on neuronal firing, that were beyond the scope of this study, our data show the feasibility of a MEA-based platform for testing drugs specifically targeting astrocytes.

Taken together, this fully human, spinal cord-specific, co-culture model with multielectrode array analyses, provides a new tool for understanding astrocyte influences on MN maturation. With the increasing use of hiPSC for modeling human diseases and for exploring therapeutic strategies, this spinal cord-specific platform offers a unique opportunity to investigate specific astrocyte/MN interactions.

## Acknowledgements

We thank Labchan Rajbhandari and Dr Venkatesan’s Lab who provided the plasma cleaning platform. Funding was provided by the Packard Center for ALS Research at Johns Hopkins. Department of Defense, W81XWH1810175. ALS Association 18-DDC-436.

The data that support the findings of this study are available from the corresponding author upon reasonable request.

---

**Supplementary figure 1:**
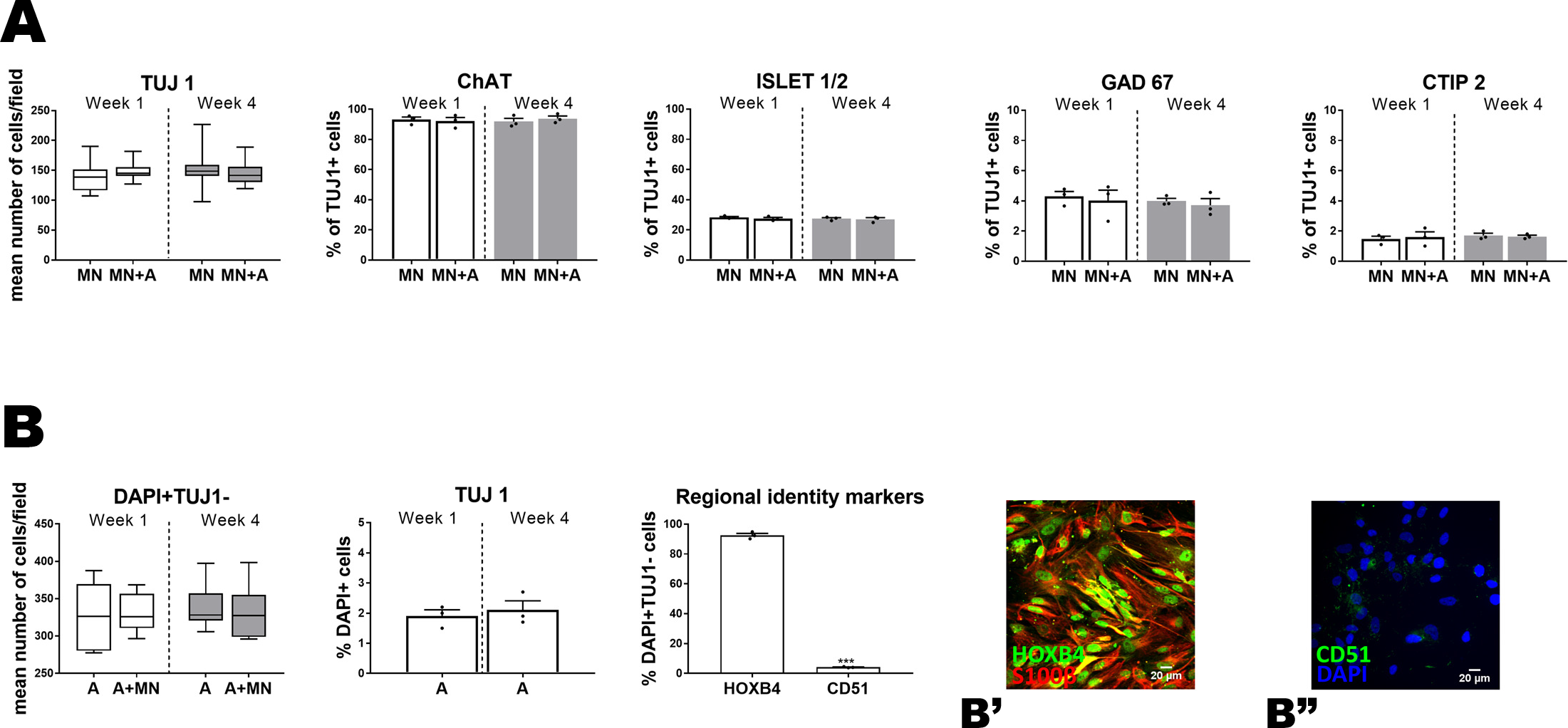
**A: Immunofluorescent quantification of neuronal markers.** The absolute number of TUJ1 immunopositive neurons (*n*=15 coverslips per culture condition and time-point) and the percentages of neuronal subtype markers (ChAT, ISLET 1/2, CTIP2, and GAD67) (*n*=3) were compared by culture condition (hiPSC-MN+A co-cultures *vs* hiPSC-MN mono-cultures) and by time of culture in vitro (week 1 *vs* week 4). **B: Immunofluorescent quantification of astrocytes markers.** The absolute number of neuronal (TUJ1^+^) and non-neuronal cells (DAPI^+^ TUJ1^−^) is shown according to co-culture conditions and by time of culture in vitro (*n*=9 coverslips per culture condition and time-point). Quantification of two regional identity markers (with representative immunostaining image, ****B’****- ****B”****), i.e. HOXB4, a markers associated with a spinal cord phenotype, and CD51, indicative of a cortical phenotype, 1 week after plating. Experimental conditions (hiPSC-MN or hiPSC-A mono-cultures vs hiPSC-MN + hiPSC-A co-cultures) are compared within each time-point after plating (*). Similarly, time-point observations (week 1 vs week 4) are compared within each condition (^). *\* or ^ p<0.05; ** or ^^ p<0.01; *** or ^^^ p<0.001.*

**Supplementary figure 2:**
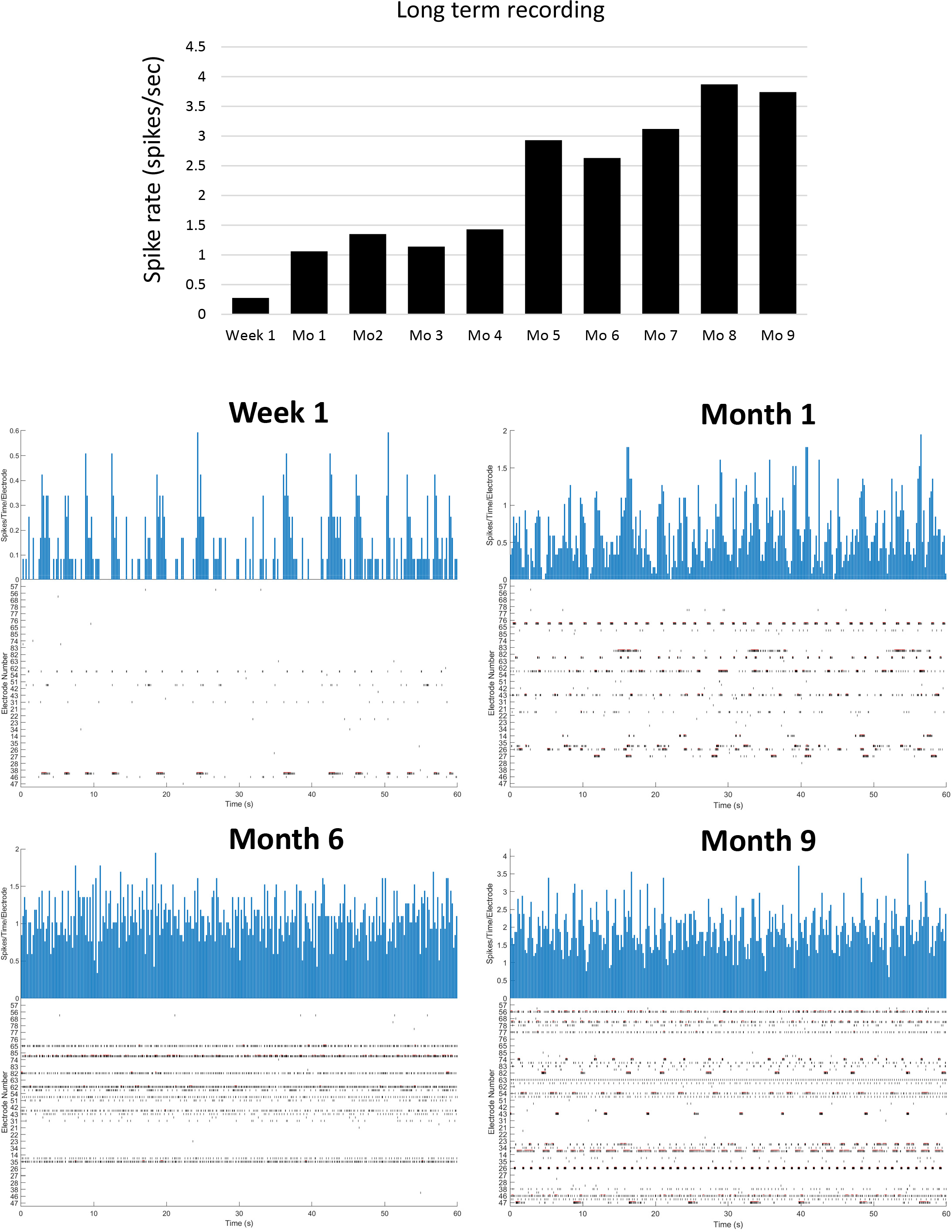
Long term MEA recording. ***Top***: hIPSC-A/hiPSC-MN co-cultures were maintained in vitro for up to 9 months and electrophysiological activity was recorded weekly. MEA recording of spike rate is reported and shows a profile with two ascending slopes and two plateau segments. ***Bottom***: Representative tracings of real-time recording at 1 week and 1, 6, and 9 months after culture.

**Supplementary figure 3:**
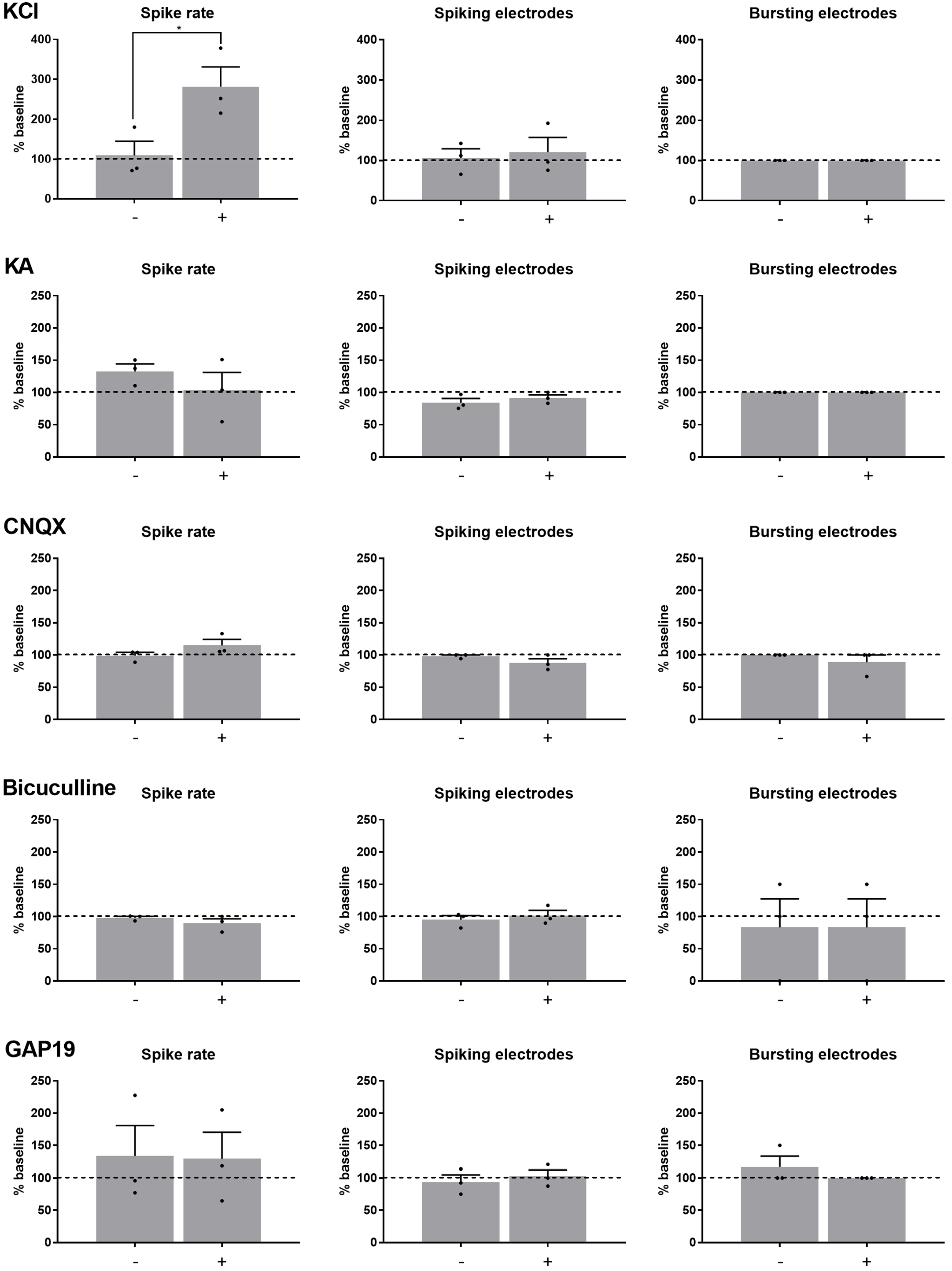
Effect of pharmacologic compounds on hiPSC-MN mono-cultures as recorded by MEA. We show the effects of the addition of KCl, a non-specific depolarizing agent, and of compounds acting on neurotransmitter receptors (KA, CNQX, bicuculline). GAP19, a drug targeting astrocyte connexin 43, was tested on cultures of neurons alone. All recordings were performed in hiPSC-MN mono-cultures 3 to 4 weeks after plating (mean of *n*=3 independent experiments). *\* p<0.05; ** p<0.01; *** p<0.001.* Abbreviations: KA, kainic acid; KCl, potassium chloride.

